# An integrated systems-biology platform for power-to-gas technology

**DOI:** 10.1101/2022.12.30.522236

**Authors:** Isabella Casini, Tim McCubbin, Sofia Esquivel-Elizondo, Guillermo G. Luque, Daria Evseeva, Christian Fink, Sebastian Beblawy, Nicholas D. Youngblut, Ludmilla Aristilde, Daniel H. Huson, Andreas Dräger, Ruth E. Ley, Esteban Marcellin, Largus T. Angenent, Bastian Molitor

## Abstract

Methanogenesis allows methanogenic archaea (methanogens) to generate cellular energy for their growth while producing methane. Hydrogenotrophic methanogens thrive on carbon dioxide and molecular hydrogen as sole carbon and energy sources. Thermophilic and hydrogenotrophic *Methanothermobacter* spp. have been recognized as robust biocatalysts for a circular carbon economy and are now applied in power-to-gas technology. Here, we generated the first manually curated genome-scale metabolic reconstruction for three *Methanothermobacter* spp‥ We investigated differences in the growth performance of three wild-type strains and one genetically engineered strain in two independent chemostat bioreactor experiments. In the first experiment, with molecular hydrogen and carbon dioxide, we found the highest methane production rate for *Methanothermobacter thermautotrophicus* ΔH, while *Methanothermobacter marburgensis* Marburg reached the highest biomass growth rate. Systems biology investigations, including implementing a pan-model that contains combined reactions from all three microbes, allowed us to perform an interspecies comparison. This comparison enabled us to identify crucial differences in formate anabolism. In the second experiment, with sodium formate, we found stable growth with an *M. thermautotrophicus* ΔH plasmid-carrying strain with similar performance parameters compared to wild-type *Methanothermobacter thermautotrophicus* Z-245. Our findings reveal that formate anabolism influences the diversion of carbon to biomass and methane with implications for biotechnological applications of *Methanothermobacter* spp. in power-to-gas technology and for chemical production.

**Graphical Abstract:** 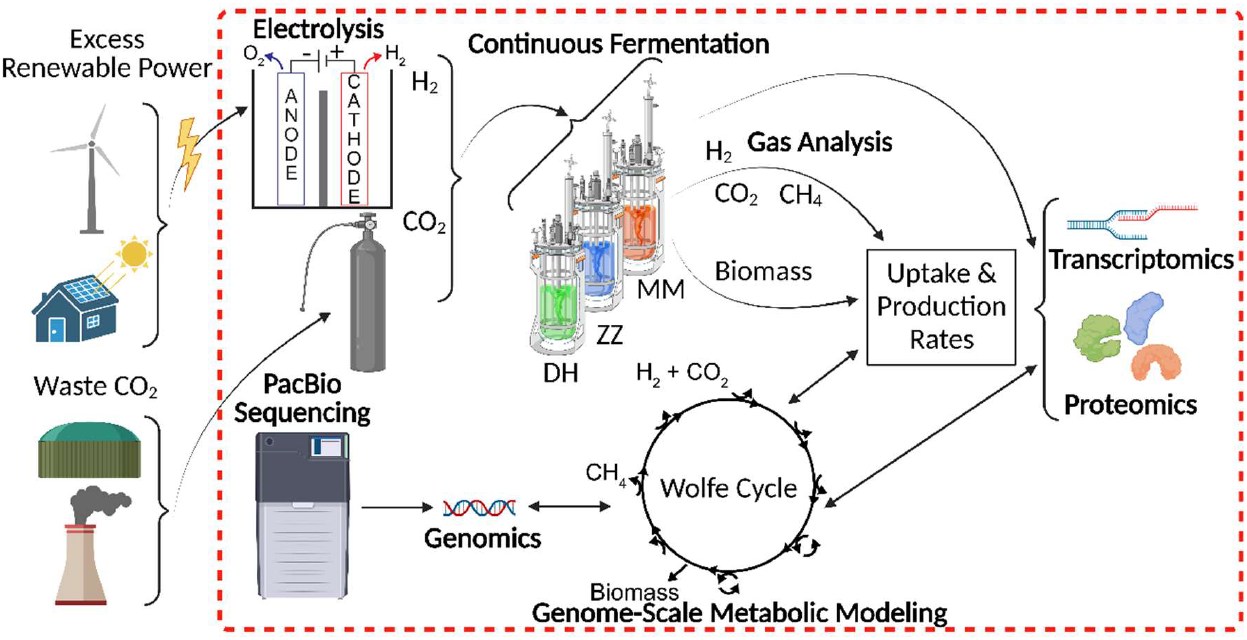

**Broader context:** Renewable energy sources (e.g., wind and solar) provide carbon-free electric power. However, their intermittency and offset between peak production and demand generate the need to store this electric power. Furthermore, these technologies alone do not satisfy the demand for carbon-based commodities. Power-to-gas technology provides a means to store intermittent renewable electric power with concomitant carbon dioxide recycling into a chemical energy carrier, such as methane, on a centralized and decentralized scale. This is particularly important to establish equitable energy strategies for *all* countries, as is highlighted by the United Nations Sustainable Development Goals. With this work, we provide an integrated systems-biology platform for *Methanothermobacter* spp. to optimize biological power-to-gas technology and formulate strategies to produce other value-added products besides methane.

## Introduction

Greenhouse-gas emissions from fossil fuels, primarily carbon dioxide, are the primary driver of anthropogenic climate change. Solutions are urgently needed to mitigate the devastating effects of greenhouse-gas emissions worldwide and decarbonize the energy and industrial sectors. Societies must reshape in a way that allows the efficient implementation of: **1)** renewable electric power to replace fossil sources for public and industrial energy demands; and **2)** carbon dioxide as feedstock instead of emission for the production of commodities within a circular carbon economy. Power-to-gas technologies are implemented to convert excess renewable electric power *via* water electrolysis into dioxygen and molecular hydrogen. In an additional methanation step, the molecular hydrogen can be combined with carbon dioxide to produce methane and water (**Equation 1**).

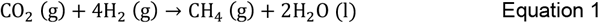

Potential carbon dioxide sources that would align with a sustainable circular carbon economy include: **1)** biogas from anaerobic digestion, which contains up to ~40% carbon dioxide;^1^ **2)** exhaust gas from non-fossil power plants such as from renewable natural gas and biomass; **3)** off-gases from other industrial processes that are difficult to be decarbonized entirely such as cement and steel production;^2–4^ and **4)** direct air capture.^5,6^ While power-to-gas can provide carbon-free molecular hydrogen, the hydrogen infrastructure is not well-established. The addition of molecular hydrogen into the existing natural gas grid is typically limited to under 10% v/v, depending on the location.^7^ However, some studies have considered higher ranges.^8–10^ Instead, methane resembles the main constituent of fossil natural gas. Methane can be injected into the natural gas grid infrastructure, which is already in place in many communities, for storage, distribution, and consumption purposes. Therefore, methane from power-to-gas has the potential to replace fossil natural gas in the natural gas grid as a decarbonization strategy and for communities to become more independent in securing energy demands.^4,11^

The methanation step in power-to-gas can be performed *via* thermo-chemical processes such as the hydrocarbon-forming Sabatier process.^12^ However, these processes typically require high temperature (> 200°C) and pressure (> 1 MPa), as well as a metal catalyst (e.g., iron, nickel, cobalt, ruthenium) that is sensitive to gas impurities.^13–15^ Alternatively, the methanation step can be performed biologically with microbes, known as hydrogenotrophic methanogenic archaea (methanogens) as biocatalysts, which naturally metabolize carbon dioxide and molecular hydrogen to produce methane (**Equation 1**).^11^,^14^,^16^ Thermophilic methanogenic species of the genus *Methanothermobacter* have been adopted for this purpose on a large scale.^17^

During the last 40 years, *Methanothermobacter thermautotrophicus* ΔH (formerly *Methanobacterium thermoautotrophicum* ΔH) and *Methanothermobacter marburgensis* Marburg (formerly *Methanobacterium thermoautotrophicum* Marburg) have served as model microbes to study hydrogenotrophic methanogenesis.^18^ This resulted in an abundance of literature on their core metabolism, which includes methanogenesis, for example, in an extensive comparative genome study.^19^ Nevertheless, more than 500 hypothetical proteins and many pathways have not been resolved.^20^ Another related species, *Methanothermobacter thermautotrophicus* Z-245 (formerly *Methanobacterium thermoformicicum* Z-245^18^), has primarily been studied for its formate metabolism^21^ and its plasmid (pFZ1).^22^ All three microbes grow on carbon dioxide and molecular hydrogen, but *M. thermautotrophicus* Z-245 can also utilize formate as a sole growth substrate (**Equation 2**) due to the presence of a catabolic formate dehydrogenase.^21^

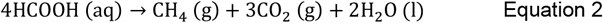

The three microbes have similar optimal growth temperatures (~65°C), pHs (~7), and no requirements for any organic compounds for growth.^23^ Further, their relatively short doubling time and growth into high biomass densities compared to other methanogens (particularly mesophilic species)^23^ promote their use for biotechnological applications. The exothermic nature of hydrogenotrophic methanogenesis renders thermophilic species especially suitable because less energy is required for cooling the bioreactor.^1^

A promising approach to predict phenotypes and identify potential bottlenecks for optimizing microbial metabolisms is through mathematical analyses of their potential metabolic networks.^24^ Genome-scale network reconstructions are knowledgebases that represent the theoretical metabolic capacities of a microbe as a network of metabolites (nodes) linked by reactions. Reactions are associated with (a set of) genes *via* the enzyme (encoded by these genes) that catalyze that respective reaction. A genome-scale metabolic model (GEM) represents the reconstruction as a stoichiometric matrix. The GEM is typically used in constraint-based modeling approaches in which steady-state conditions and conservation of energy and mass are assumed. These mathematical models enable virtual experiments, including investigating gene deletions and insertions. GEMs comprehensively organize the knowledge about microbes and systematize our understanding of how they live and grow. During the last 20 years, thousands of reconstructions and GEMs have been assembled across all three domains of life; however, archaea remain underrepresented. Only 127 of 6239 (2%) are archaeal GEMs.^25^ Of those, only ten models have been manually curated, which is a procedure that leads to higher quality models.^25^ Many tools have been developed to accelerate model-building, mainly through automation^26^ – a process that often relies on genome annotations, biochemical data, and other models from databases, which are more limited and less specific for archaea.^27^

The applications of GEMs and related models have been extensively reviewed.^28^ Besides predicting phenotypes and bottlenecks, GEMs also provide a backbone for integrating cultivation data, omics data (genomics, transcriptomics, proteomics, metabolomics, and fluxomics), kinetics data, and thermodynamics data through various methods from flux balance analysis to machine learning.^28^ Further, the reconstruction of pan-models or pan-GEMs enables: **1)** the construction of GEMs for new microbes, especially non-model microbes;^29^ **2)** the comparative genomic-to-phenotypic study of closely related microbes;^29,30^ and **3)** the analysis of the evolution of metabolism of the selected species (if constructed using orthologous genes).^31^ Few studies collected samples of *Methanothermobacter* spp. for fermentation analyses during steady-state from chemostat bioreactors.^21,32–34^ Only limited studies have looked at the entire transcriptome or total proteome of *M. thermautotrophicus* ΔH^35,36^ or *M. marburgensis* Marburg.^37^ These studies investigated the effect of varying cultivation conditions on the transcriptome^35^ and proteome.^36,37^ However, no published omics data are available for *M. thermautotrophicus* Z-245. Further, no study compared the methanogens using pan-transcriptomics and pan-proteomics (in general, few studies compare directly across closely related species^38,39^).

To assess the differences and potential advantages for bioprocessing between the three different model *Methanothermobacter* spp., we: **1)** (re)sequenced the genomes (*M. thermautotrophicus* Z-245 genome sequence was not available before); **2)** performed a genome comparison (*i.e*., created a pan-genome by identifying homologous genes across the three microbes); **3)** reconstructed a high-quality manually-curated pan-model and GEMs; **4)** operated two experiments with quadruplicate chemostat bioreactors to achieve steady-state conditions; **5)** collected transcriptomics and proteomics data sets from these bioreactors; and **6)** integrated these data sets into the GEMs. We compared the three wild-type microbes and a plasmid-carrying *M. thermautotrophicus* ΔH strain (*M. thermautotrophicus* ΔH pMVS1111A:P_*hmtB*_-*fdh*_Z-245_) that is capable of utilizing formate as a growth substrate.^40^ With these extensive data sets, we demonstrated how a systems-biology approach can be applied for interspecies comparisons while providing a rich collection of multi-level data that will be a valuable resource for the scientific community, for example, to help fill the knowledge gaps in archaeal metabolism.

## Results and Discussion

### Updated genome sequences for *Methanothermobacter* spp. provide the basis for high-quality genome-scale metabolic reconstructions

We conducted independent *de-novo* sequencing and genome assembly for *M. thermautotrophicus* ΔH, *M. thermautotrophicus* Z-245, and *M. marburgensis* Marburg with long-read sequencing technology (**Table 1; Table S1 and Section 1.2-1.3, ESI†** [detailed descriptions of the materials and methods are provided in Section 1, **ESI†**]). This procedure provided high-quality genome sequences (first release of the *M. thermautotrophicus* Z-245 genome sequence), including the species-specific methylation pattern of the three microbes (**Text S1 and Table S2, ESI†**). In our genome annotation, we assigned COG functional annotations to 1501/1796, 1505/1804, and 1440/1730 genes for *M. thermautotrophicus* ΔH, *M. thermautotrophicus* Z-245, and *M. marburgensis* Marburg, respectively (**Table S3, ESI†**). We identified genes that were: **1)** previously predicted but not identified in our new sequences for *M. thermautotrophicus* ΔH and *M. marburgensis* Marburg; and **2)** identified but not previously predicted (**Table S4, ESI†**), which we considered for the GEM reconstruction described below. For example, we did not identify a predicted ribonucleoside-diphosphate reductase in *M. marburgensis* Marburg (MTBMA_c10330) in our new genome sequence. These differences were likely due to advances in sequencing methods and algorithms for assembly and annotation during the last few decades (*M. thermautotrophicus* ΔH was sequenced in 1997^41^ and *M. marburgensis* Marburg in 2010^42^). For sequencing, we used strains directly from the DSMZ without excessive subcultivation (**Section 1.1, ESI†**). However, we cannot exclude differences due to lab-culture adaptation.

**Table 1.** Summary of sequenced genomes.

| Microbe | Plasmid | Length<br>(bps) | GC % | Gene<br>Count | CDS | Previous Sequence<br>Accessions | Difference<br>with Old<br>Sequence | Similarities<br>with Old<br>Sequence |
| --- | --- | --- | --- | --- | --- | --- | --- | --- |
| *M. thermautotrophicus*<br>ΔH | N/A | 1,751,429 | 49.56 | 1844 | 1796 | NC_000916.1<sup>41</sup> | 448 SN<sup>1</sup><br>247 IB<sup>2</sup><br>91 O2N<sup>3</sup><br>92 N2O<sup>4</sup> | 1733 CDS |
| *M. thermautotrophicus*<br>Z-245 | pFZ1 | 1,758,798<br>11,015<sup>5</sup> | 49.46<br>42.51<sup>5</sup> | 1840<br>12<sup>5</sup> | 1792<br>12<sup>5</sup> | GCA_013330715.1<sup>6</sup><br>80 | N/A | N/A |
| *M. marburgensis*<br>Marburg | pM2001<br>(pMTBMA4) | 1,634,705<br>4,441<sup>5</sup> | 48.65<br>45.40<sup>5</sup> | 1774<br>5<sup>5</sup> | 1725<br>5<sup>5</sup> | NC_014408.1<sup>42</sup> | 68 SN<br>30 IB<br>25 O2N<br>66 N2O | 1675 CDS |

<sup>1</sup> SN, single nucleotides
<sup>2</sup> IB, indel bases
<sup>3</sup> O2N, old genes that do not map to any new genes, **Table S4, ESI†**
<sup>4</sup> N2O, new genes that do not map to any old genes, **Table S4, ESI†**
<sup>5</sup> indicates values for the plasmid
<sup>6</sup> sequence was from a metagenome and the assembly only to scaffold level

Additionally, the gene annotation style was slightly altered. Specifically, stop codons no longer inevitably resulted in the prediction to terminate a gene. For example, the two homologs of adenosine monophosphate (AMP)-forming acetyl-CoA synthetase (ACS), which catalyze the formation of acetyl-CoA from acetate using ATP, were both split into two open reading frames (ORF) in the old *M. thermautotrophicus* ΔH genome annotation (MTH217-MTH216 and MTH1603-MTH1604) (**Table 2**). In our new annotation, both sets of genes mapped to only one ORF (ISG35_00975 and ISG35_07685). In the first set, there remains an in-frame stop codon located in the middle of the gene (at amino acid number 559), which was noted with the comment “internal stop” in the GenBank file (in total, four genes in the genome have this note). This stop codon likely renders this ACS non-functional in *M. thermautotrophicus* ΔH.^43^ However, in the second set, the new gene annotation (ISG35_07685) did not contain an in-frame stop codon anymore, confirming the error in the old genome sequence.^43^ The second ACS could be characterized before.^43^ However, our omics data revealed that it was transcribed, translated, and thus, assigned to a reaction in the pan-model (**Data S1, ESI†**). In addition, a putative acetate transporter was annotated in both the previously reported (MTH215) and new (ISG35_07680) sequence, and we found this gene to be transcribed and translated (**Table 2**). It remains to be elucidated experimentally what implications these findings have for the acetate metabolism in *M. thermautotrophicus* ΔH. Overall, the genome sequences with the updated annotations provided a high-quality basis for the construction of our GEMs.

**Table 2.**
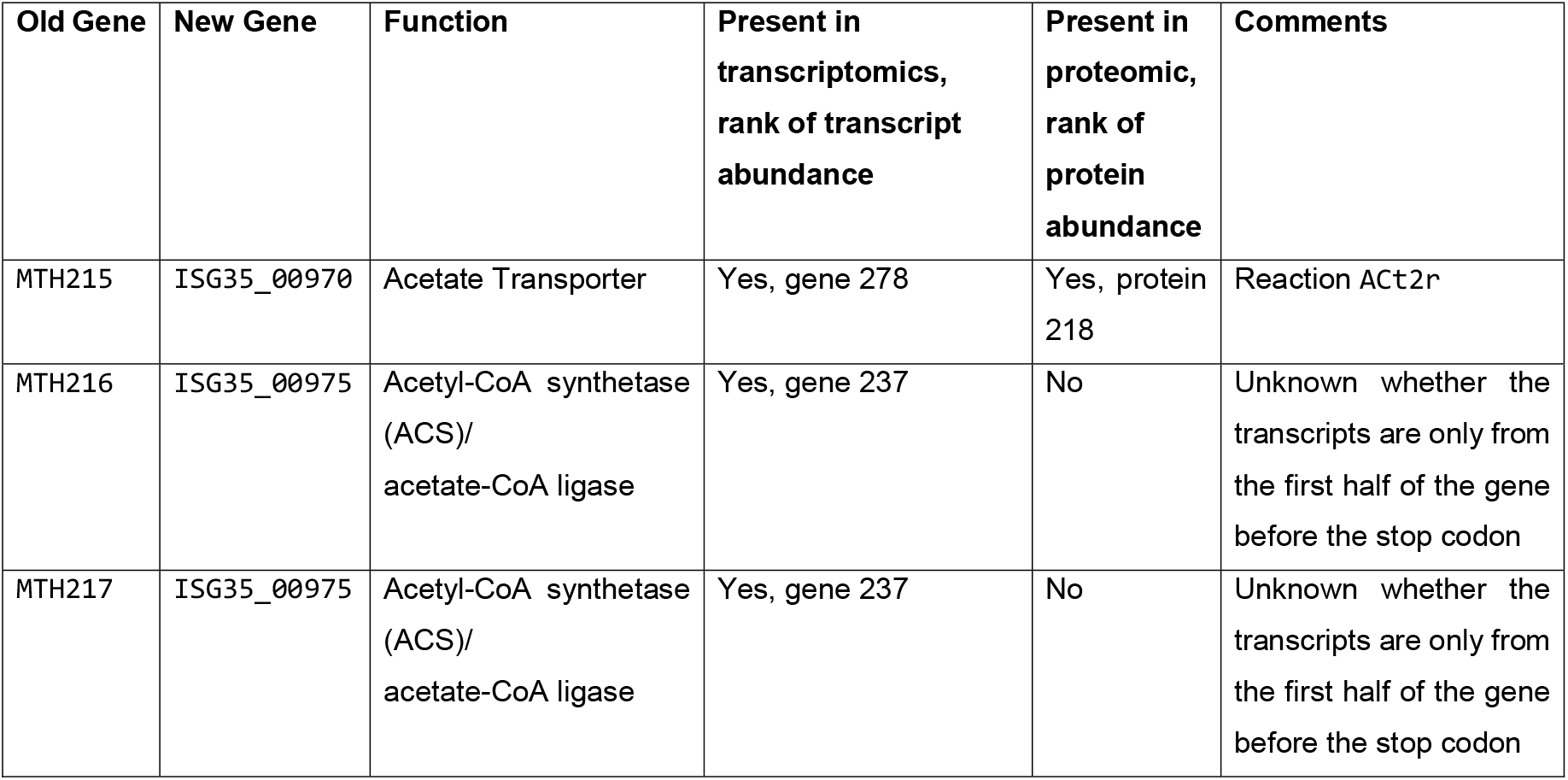
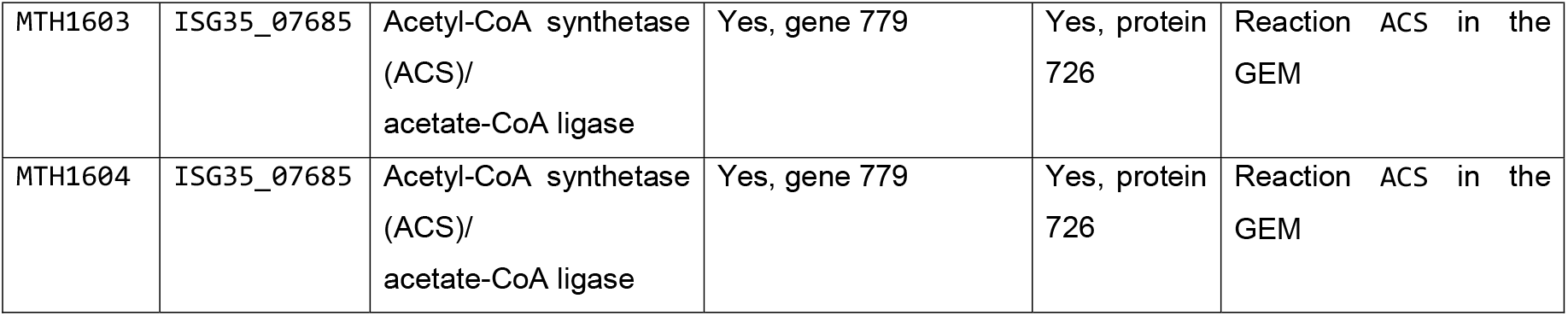
The comparison of the acetate transporter and acetyl-CoA synthetase/acetate-CoA ligase genes in the old and new genome sequences and their presence in the transcriptomics and proteomics data of *Methanothermobacter thermautotrophicus* ΔH. The rank of transcript and protein abundance is also listed.

### Our pan-model results in a high genome coverage compared to current methanogen reconstructions

To streamline a comparison between our three microbes, we constructed a pan-genome-scale metabolic reconstruction (pan-model) that integrates the metabolic capabilities of all three microbes. This pan-model consists of 618 reactions (including 46 exchange reactions, 56 transport reactions, and seven biomass-associated reactions, with eleven transport and eleven exchange reactions that act as pseudo reactions for orphan metabolites to refer to compounds that are either only produced or consumed), 555 metabolites, and 545 genes (**Data S2, ESI†**). The pan-model reflects 29.3% (526/1796, *M. thermautotrophicus* ΔH), 28.8% (520/1804, *M. thermautotrophicus* Z-245), and 29.4% (509/1730, *M. marburgensis* Marburg) of the protein-coding genes. This coverage for archaea, specifically methanogens, is comparable to published reconstructions, for which the average is 20% and the highest 29%.^44^ Based on the pan-model, we derived strain-specific GEMs for each of the three microbes (*i*MTD22IC, *i*MTZ22IC, *i*MMM22IC), which we used for further modeling as described below (**Data S2, ESI†**). We followed recent recommendations for best practices, including for the GEM nomenclature.^45^ We tested the GEMs in MEMOTE, which is a tool that provides a comparable score about the quality of a model, and each scored 85% (**Data S2, ESI†**).^46^ Older models tend to score lower; however, even a model that was built using the relatively new automated reconstruction tool, CarveME,^47^ only achieved 33%.^24^ Further, we produced metabolic pathway maps to visualize the GEMs with Escher (**Data S2, ESI†)**.^48^ In this study, we utilized the pan-model and GEMs to explain observed differences between the three microbes.

### *M. thermautotrophicus* ΔH has a significantly higher methane-production rate compared to *M. thermautotrophicus* Z-245 and *M. marburgensis* Marburg

To compare the metabolism of the three microbes and to produce statistically relevant data to constrain the GEMs, we performed quadruplicate chemostat bioreactors (*N*=4) with the three microbes in pure culture under the same growth conditions in a single experimental run. For *M. thermautotrophicus* Z-245, we discarded one replicate (*N*=3) due to a pump malfunction in this bioreactor and wash-out of the cells (**Section 1.8, ESI†**). We performed PCR analyses with strain-specific primer pairs, which excluded cross-contamination between the bioreactors (**Figure S1 and Section 1.9, ESI†**). Under steady-state growth conditions, *M. thermautotrophicus* ΔH had consumption rates for molecular hydrogen of 234.36 ± 40.76 mmol/gCDW/h and carbon dioxide of 58.18 ± 11.49 mmol/g_CDW_/h, with a production of 52.50 ± 8.53 mmol/gCDW/h methane and 0.03 ± 0.01 gCDW biomass (**Figure 1A, B; Table S5, ESI†**). This was a significantly higher consumption rate for molecular hydrogen (1.59-fold and 1.72-fold) and carbon dioxide (1.69-fold and 1.72-fold), as well as production rate for methane (1.62-fold and 1.82-fold), compared to both *M. thermautotrophicus* Z-245 and *M. marburgensis* Marburg (**Figure 1A, B; Data S3, ESI†**). However, the biomass concentration of 0.04 ± 0.01 gCDW was significantly (1.36-fold) higher for *M. marburgensis* Marburg than for the other two microbes (**Figure 1A, B; Data S3, ESI†**). This divergence resulted in differences in the normalized distribution of products, with the highest methane-to-biomass ratio of 96.4 ± 0.57 for *M. thermautotrophicus* ΔH and the lowest of 93.5 ± 0.98 for *M. marburgensis* Marburg under our experimental conditions (**Figure 1C**).

**Figure 1.**
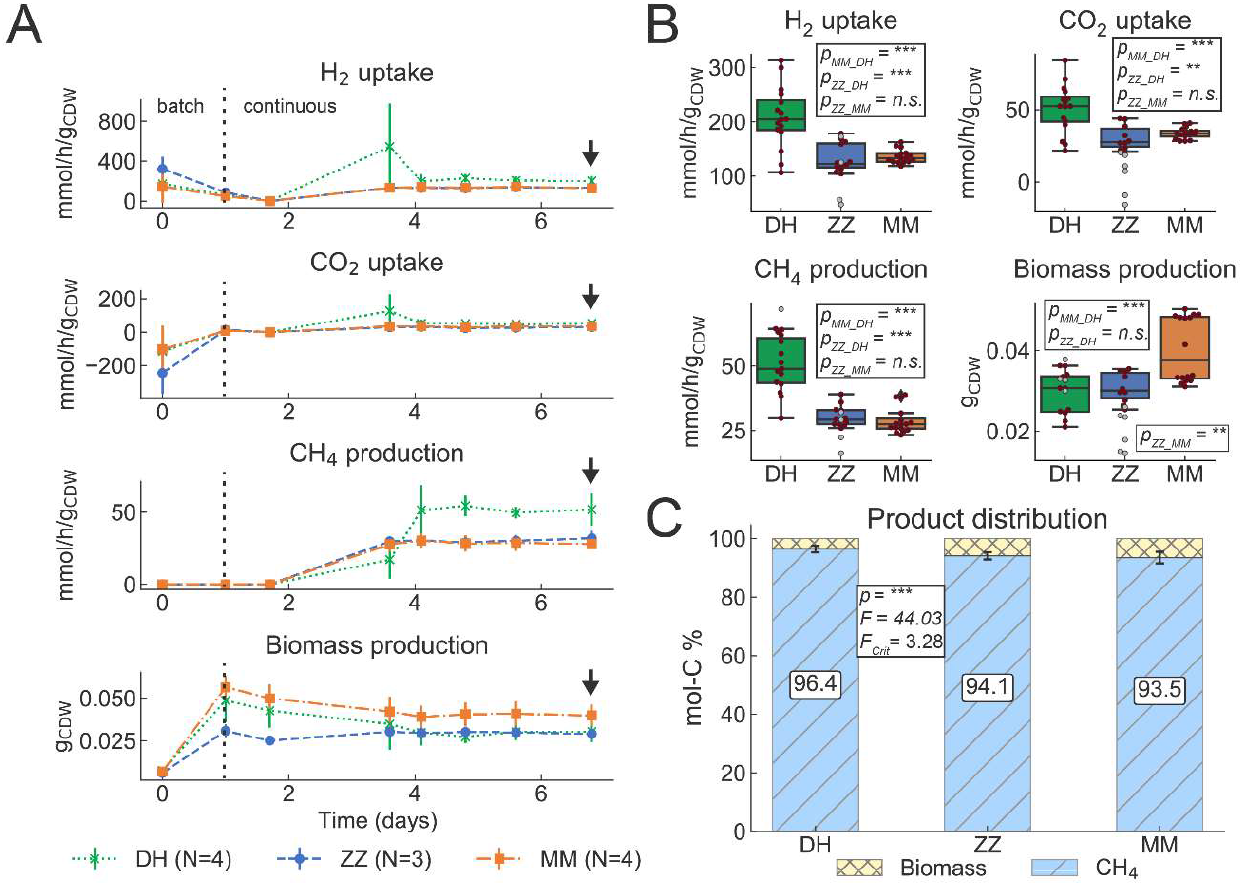
First experiment fermentation data from chemostat bioreactors with *M. thermautotrophicus* ΔH (DH), *M. thermautotrophicus* Z-245 (ZZ), and *M. marburgensis* Marburg (MM). **A)** Gas consumption (H_2_ and CO_2_ uptake), and CH_4_ and biomass production data from quadruplicate (DH and MM) and triplicate (ZZ) bioreactors for the fermentation period of 7 days. Data for further analyses (transcriptomics, proteomics) were taken on day seven, as indicated by arrows. **B)** Average gas consumption (H_2_ and CO_2_ uptake) and CH_4_ and biomass production data during a steady-state period (days 4 to 7). For statistical analysis in pair-wise comparisons with *t*-test, data points without suspected gross measurement error (red circles) were included, data points with suspected gross measurement error (gray circles) were excluded (**Section 1.13, ESI†**). **C)** Average normalized product distribution, including statistical analysis by ANOVA (**Section 1.14, ESI†**). DH, *M*. *thermautotrophicus* ΔH; ZZ, *M. thermautotrophicus* Z-245; MM, *M. marburgensis* Marburg; ***, *p* < 0.0001; **, *p* < 0.001; n.s., not significant (p > 0.05); F, F value; Fcrit, F critical value.

Based on our findings, for industrially relevant power-to-gas systems, the clear choice between these three methanogens would be *M. thermautotrophicus* ΔH due to its superior kinetics and methane specificity. However, it is essential to consider the fermentation conditions of our study. We applied a volume gas per volume bioreactor per minute (vvm) of 0.08, while others applied a much higher vvm of up to 2.01.^17^ Whether the superior behavior of *M. thermautotrophicus* ΔH would hold at the higher vvm was not evaluated here, and should be considered. Here, we did not maximize methane production rates or achieve high methane partial pressures in the product gas. Instead, we produced replicate data sets under identical experimental conditions for the three microbes.

### Interspecies comparison of multi-level omics under steady-state growth conditions reveals different gene-expression patterns

For an in-depth assessment of the bioreactor experiments, we performed transcriptomics and proteomics analyses during steady-state for all replicate bioreactors (**Data S1, ESI†**). We achieved high reproducibility of the replicates for both transcriptomics and proteomics and high coverage of the transcriptome (94.18-99.77%) and proteome (78.29-79.82%) with each microbe (**Table 3A**). The high coverage of both the transcriptome and the proteome indicates that the density of coding sequences in the genomes of the microbes is further reflected at the transcript and protein levels. The numbers of identified proteins in our study are similar to those previously found for *M. thermautotrophicus* ΔH by Liu, *et al*.^36^

**Table 3.**
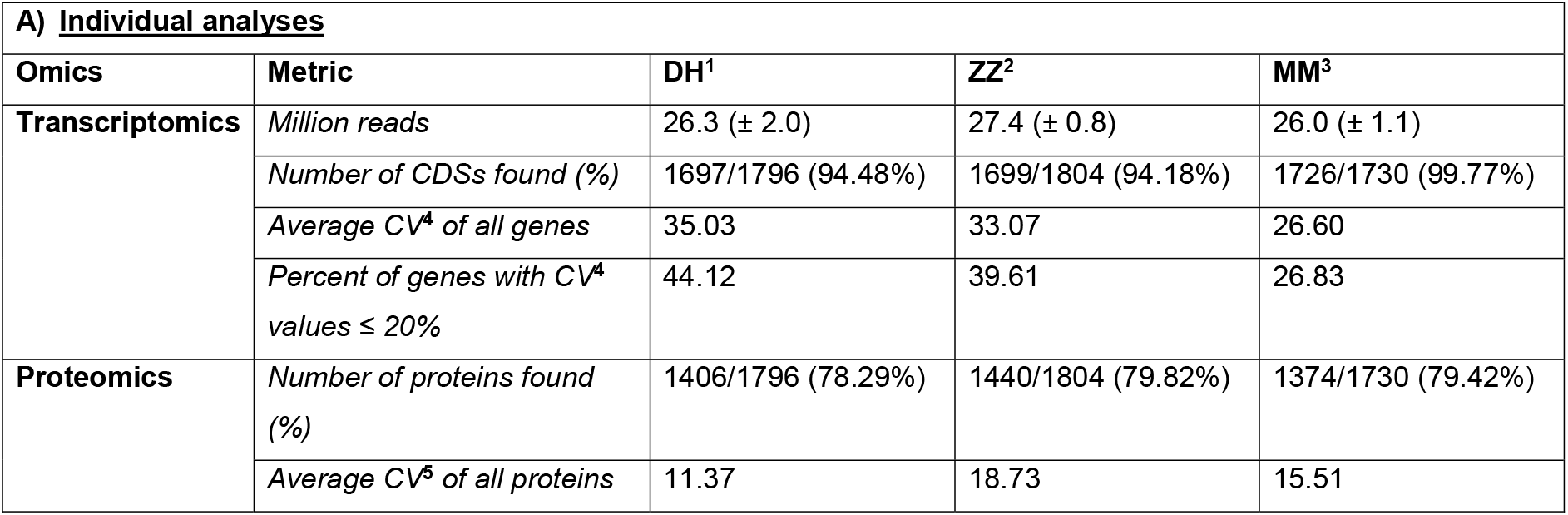
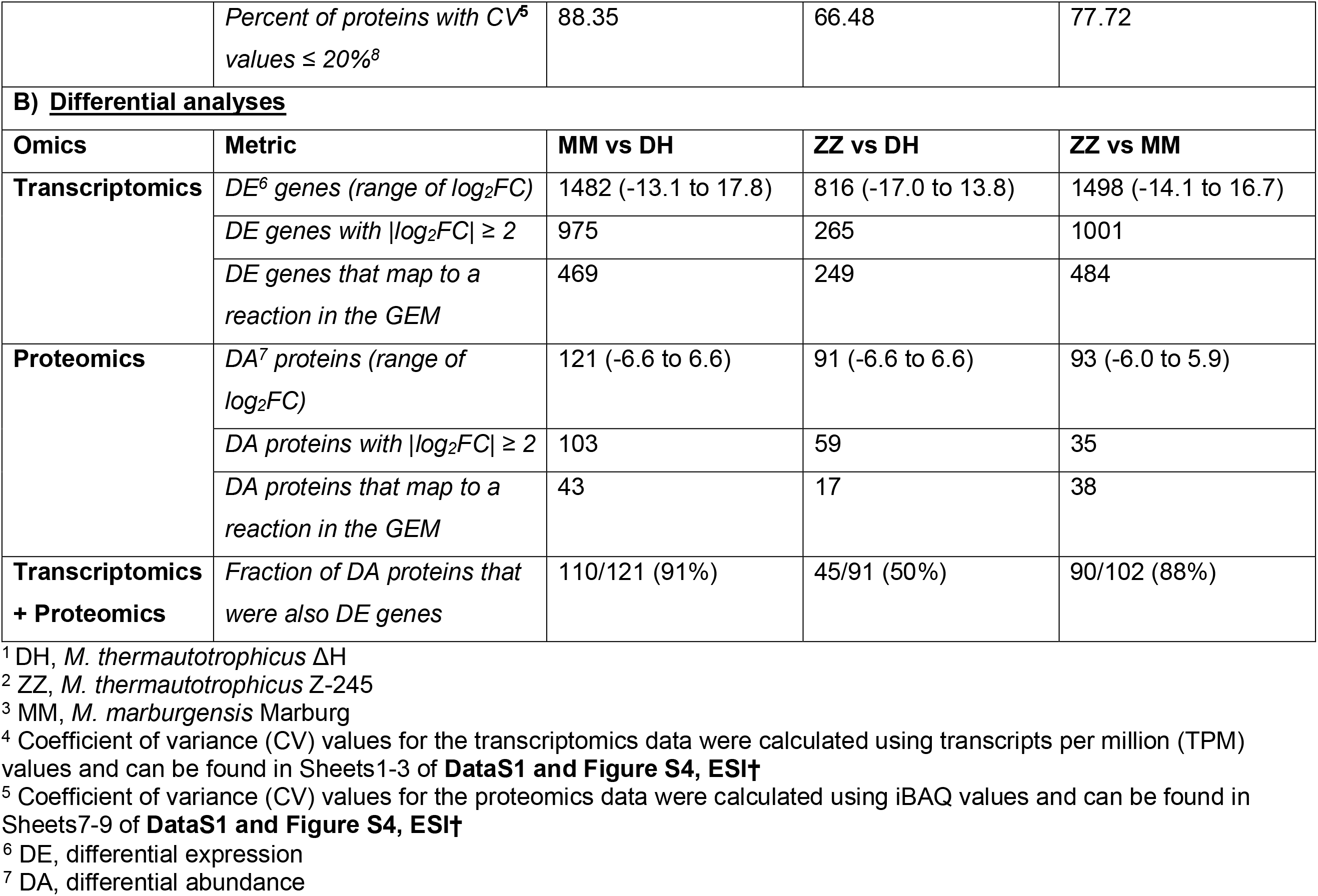
Metrics of the transcriptomics and proteomics for the individual analyses and the differential expression and differential abundance analyses for the three microbes. Statistical significance was determined by an adjusted P-value ≤ 0.05.

To compare the omics data sets between microbes, we merged the underlying genomes into pan-transcriptome and pan-proteome databases (creating reference genes, termed “gene groups,” **Data S1, ESI†, Section 1.22, ESI†**). Merging allowed us to perform differential expression analyses for the pair-wise comparison of *M. marburgensis* Marburg versus *M. thermautotrophicus* ΔH (MM/DH), *M. thermautotrophicus* Z-245 versus *M. thermautotrophicus* ΔH (ZZ/DH), and *M. thermautotrophicus* Z-245 versus *M. marburgensis* Marburg (ZZ/MM). We found many differentially expressed genes in the pan-transcriptome but much fewer differentially abundant proteins in the pan-proteome comparison, and in both omics sets, fewer differences between *M. thermautotrophicus* ΔH and *M. thermautotrophicus* Z-245 than to *M. marburgensis* Marburg (**Table 3B; Figure 2C-G**; **Table S6 and Data S1, ESI†**). In addition, the 100 most highly expressed genes on the transcript level share few overlaps (**Figure 2A**), while we found a high overlap of the 100 most abundant proteins between microbes (**Figure 2B**). This finding was also reflected in the pathway distribution when mapping these highly expressed genes or abundant proteins to the GEMs (**Table S7 and Table S8, ESI†**).

**Figure 2.**
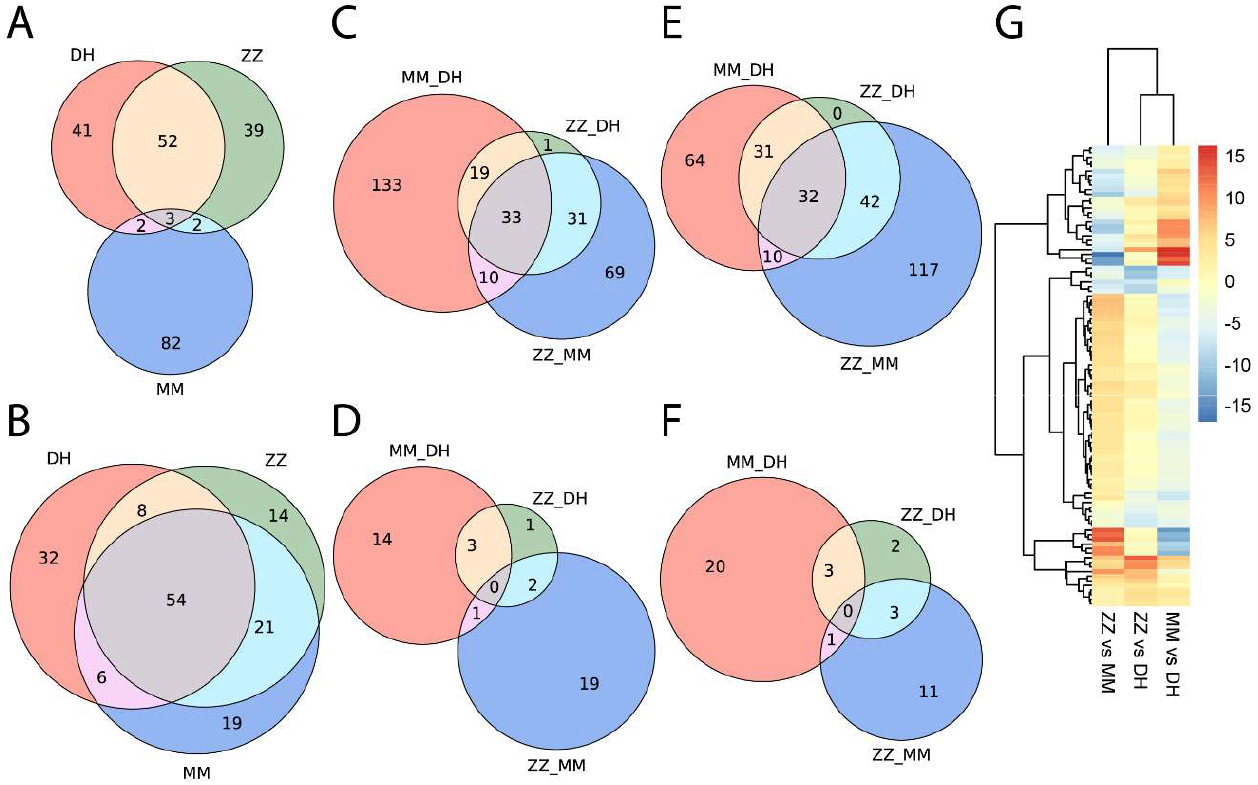
Overview of transcriptomics and proteomics interspecies comparison. **A)** Comparison of the top 100 transcribed genes (by average number of transcripts per million). **B)** Comparison of the top 100 abundant proteins (by average iBAQ value). Transcripts/proteins that do not map to a gene group are counted only once collectively; hence there are less than 100 genes total in **A** and **B**. **C)** Upregulated transcripts; **D)** more abundant proteins; **E)** downregulated transcripts; **F)** less abundant proteins, respectively, from the pair-wise differential expression analysis that map to reactions in the GEMs. **G)** Heatmap of log_2_FC in transcriptomes top 100. DH, *M. thermautotrophicus* ΔH; ZZ, *M. thermautotrophicus* Z-245; MM, *M. marburgensis* Marburg.

We did not find a correlation between the differential expression analysis of the transcriptome and proteome (**Figure S2, ESI†**), which is an observation made before with other microbes.^49^ A low correlation between differentially expressed genes and differentially abundant proteins may be explained with biological and experimental reasons.^50–53^ Transcriptional and post-transcriptional regulation and processes determine protein abundances at steady-state, specifically, rates of: **1)** transcription; **2)** translation; **3)** mRNA degradation; and **4)** protein degradation.^50,51,54,55^ The same protein abundance can be reached in countless ways with varying rate combinations, even by just varying transcription and translation rates (*e.g*., low mRNA abundance with high translation rates *versus* high mRNA abundance with low translation rates).^55^ Transcripts (mRNA) have shorter half-lives (faster degradation rates) than proteins and, thus, are generally considered less stable.^52–54^ Distinct proteins have differing half-lives that are not necessarily correlated to the quantity of mRNA (e.g., low-abundance proteins may be detected as such due to quick degradation regardless of mRNA concentrations).^56^ In addition, experimental biases are known.^50^ Assuming that the data quality of each method was high (discussion on coefficients of variance [CVs] in next section), we hypothesized that regulation patterns of each methanogen may have evolved differently but that the outcome on proteome level was very similar to maximize growth under thermodynamically limited conditions. We continued to investigate this here, especially to explain the highest methane-to-biomass ratio for *M. thermautotrophicus* ΔH, which is pertinent for bioprocessing.

### Differential expression of methanogenesis pathway genes does not mirror significant differences in growth behavior

Although the conditions in the bioreactor experiments were identical for all three closely related species, the growth behavior and overall transcriptome and proteome composition between the microbes revealed significant differences. To gain deeper insight into possible explanations for these differences, we took a closer look at methanogenesis as the core metabolic feature of methanogens. The differential transcriptomic analysis generally indicated that for *M. marburgensis* Marburg, genes of the Wolfe cycle (*i.e*., methanogenesis)^57^ were upregulated compared to the other two microbes (**Figure 3; Figure S3 and Data S1, ESI†**). There were, however, some genes that were downregulated, including F_420_-non-reducing Ni-Fe hydrogenase (Mvh), molybdenum-dependent formylmethanofuran dehydrogenase (Fmd), and several subunits of the F_420_-reducing Ni-Fe hydrogenase (Frh), methyl-tetrahydromethanopterin (H_4_MPT) coenzyme M methyltransferase (Mtr), and methyl-coenzyme M reductase isoenzyme II (Mrt). Nonetheless, these differences on transcriptome level did not translate to the proteome level to the same degree, and only some of the proteins were differentially abundant between the three microbes, as outlined in the previous section (**Figure 3**; **Data S1, ESI†**).

**Figure 3.**
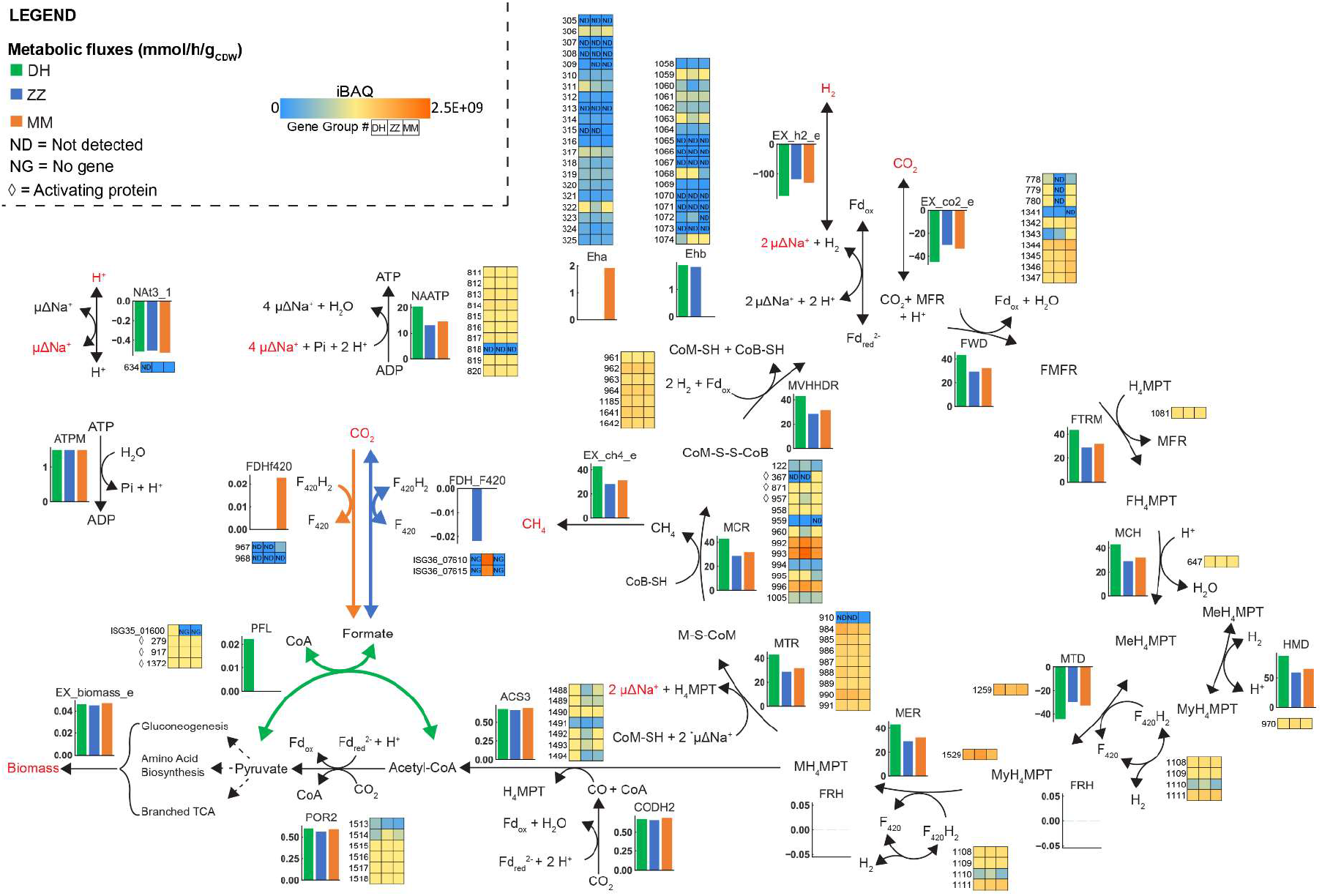
Wolfe Cycle adapted from Thauer^57^ with other reactions involved in the energy metabolism. Fluxes are from the proteomics reduced model (using iBAQ values), constrained with experimental data that was adjusted for gross measurement error. Gene group is used as ID for the omics. For the PFL reaction, only *M. thermautotrophicus* ΔH has the gene; thus, the gene ID is used (ISG35_01600). For the FDH_F420, only *M. thermautotrophicus* Z-245 has the formate dehydrogenase cassette; thus, the gene IDs are used (ISG36_07610 and ISG36_07615). Red text refers to metabolites that are exchanged across the membrane. **Microbes**: DH, *M. thermautotrophicus* ΔH; MM, *M. marburgensis* Marburg; ZZ, *M. thermautotrophicus* Z-245. **Compounds**: CH_4_, methane; CO, carbon monoxide (carbonyl group); CoA, Coenzyme A; CoB, coenzyme B; CoM, coenzyme M; CoM-S-S-CoB, CoM-CoB heterodisulfide; CO_2_, carbon dioxide; F, formyl; Fdox/rd, ferredoxin oxidized/reduced; H_2_, hydrogen; H^+^, proton; H_4_MPT, tetrahydromethanopterin; M, methyl; Me, methenyl; MFR, methanofuran; My, methylene; Na^+^, sodium ion. **Reactions/Enzymes:** ATPM, ATP maintenance (pseudo reaction); CODHr2, CO dehydrogenase/acetyl-CoA synthase; Eha/Ehb, energy converting hydrogenases; EX_biomass_e, biomass exchange (pseudo reaction); EX_ch4_e, CH_4_ exchange (pseudo reaction); EX_co2_e, CO_2_ exchange (pseudo reaction); EX_h2_e, H_2_ exchange (pseudo reaction); FDHf420, F_420_-dependent formate dehydrogenase; FDH_F_420_, F_420_-dependent formate dehydrogenase cassette; FRH, F_420_-reducing hydrogenase; FTRM, FMFR/H_4_MPT formyltransferase; FWD, FMFR dehydrogenase (tungsten- and molybdenum-dependent isozymes); HMD, MeH_4_MPT hydrogenase; MCH, MeH_4_MPT cyclohydrolase; MCR, MCoM reductase (I and II); MER, MyH_4_MPT reductase; MTD, MyH_4_MPT dehydrogenase; MTR, MH_4_MPT/CoM methyltransferase; MVHHDR, F_420_-non-reducing hydrogenase with the heterodisulfide reductase; NAATP, ATP synthase; Nat3_1, Na^+^/H^+^ antiporter; PFL, pyruvate formate-lyase; POR2, pyruvate synthase. **Other:** ⋄, activating protein; ND, not detected; NG, no gene.

To confirm the overall quality of the omics results, we analyzed the CVs for the transcriptomics and proteomics data (**Table 3; Figure S4, ESI†**). We found a larger variation in the CVs of the transcriptomics data compared to the CVs of the proteomics data. This was consistent with more differentially expressed genes than differentially abundant proteins (**Table 3; Figure S4, ESI†**). Overall we found: **1)** a high number of genes and proteins identified; **2)** CVs that are especially good for the proteomics data; **3)** a larger variance in the differential expression of the transcriptome compared to the proteome; and **4)** a large number of differentially abundant proteins that we also identified as differentially expressed genes. This analysis supported high-quality omics data and, in particular, revealed a more plastic transcriptome and a more stable proteome (**Table 3**).

Further, the relative abundances of the different protein subunits in each protein complex and the relative abundance of proteins involved in methanogenesis compared to the entire proteome were found to be similar between the three microbes (**Table S9 and Data S1, ESI†**). The most abundant methanogenesis-related proteins were Mtr, methyl-coenzyme M reductase isoenzyme I (Mcr), and tungsten-dependent formyl-MFR dehydrogenase (Fwd) for *M. thermautotrophicus* ΔH, *M. thermautotrophicus* Z-245, and *M. marburgensis* Marburg, respectively. All three microbes had significantly higher abundances of Mcr to Mrt (P-value = 0.0019), Fwd to Fmd (P-value = 0.0221), and F_420_-dependent methylene-H_4_MPT dehydrogenase (Mtd) to H_2_-forming methylene-H_4_MPT dehydrogenase (Hmd) (P-value = 0.0013) (**Table S9, ESI†**). In addition, all three microbes had similar ratios of the energy-converting hydrogenases Eha and Ehb on proteome level (P-value = 0.7951) (**Table S10, ESI†**). Overall, differences in the abundance of methanogenesis-related proteins did not explain differences in methane production between the microbes.

As mentioned before, *M. marburgensis* Marburg had a higher biomass production rate (0.01 ± 0.002 g_CDW_/L/h) compared to *M. thermautotrophicus* ΔH (1.40-fold) and *M. thermautotrophicus* Z-245 (1.39-fold). This was not affected by an air intrusion (that led to higher dioxygen levels) into one of the *M. marburgensis* Marburg bioreactors (**Figure 1B-C**; **Figure S5, ESI†**), which we confirmed by the uniformity of the CVs between the microbes (average ± standard deviation, median): *M. thermautotrophicus* ΔH (13.18 ± 12.38 %, 10.05 %); *M. thermautotrophicus* Z-245 (21.81 ± 15.66 %, 18.53 %); *M. marburgensis* Marburg (18.06 ± 15.34 %, 14.23 %). While this dioxygen intrusion had no detectable impact on the global CVs, the intrusion led to an elevated oxidative-stress response, which we could detect in the transcriptome and proteome of *M. marburgensis* Marburg (**Text S2, ESI†)**.

Because we did not find an explanation for the different growth behaviors thus far, we considered next the first few reactions that branch off from methanogenesis towards biomass growth (e.g., ACS, CODH/CODHr2, POR2, HMGCOAS, PC). However, we still did not find any clear patterns that may explain the different growth behaviors (**Data S1, ESI†**). Thus, we looked at metabolic functions that are related to biomass growth downstream in the metabolism. We used the GEMs and found that the higher biomass production rates in *M. marburgensis* Marburg were partially supported when considering the expression of metabolic genes that are related to biomass growth. Specifically, compared to *M. thermautotrophicus* ΔH, 199 genes were upregulated, and 132 genes were downregulated, while compared to *M. thermautotrophicus* Z-245, 198 genes were upregulated, and 163 genes were downregulated in *M. marburgensis* Marburg, respectively (**Table S11, ESI†**). The pathways with the most differentially expressed genes (up and down) include: **1)** purine biosynthesis; **2)** methanogenesis cofactor biosynthesis pathways (adenosylcobalamin and methanopterin); and **3)** central carbon metabolism (TCA, gluconeogenesis, and mixed sugar metabolism) (**Table S11, ESI†**). Overall, the omics data alone did not provide an explanation for the observed differences in growth behavior. Thus, we integrated the omics and fermentation data into the GEMs to better understand these differences between the three microbes.

### Integration of experimental data into GEMs results in feasible solutions for flux balance analyses and reveals differences in the generation of formate for biomass growth

With the steady-state fermentation data from the bioreactor experiment (carbon dioxide and molecular hydrogen consumption rates and methane and biomass production rates), we constrained the GEMs in flux balance analyses (**Section 1.25, ESI†**). Importantly, we corrected the consumption and production rates for experimental errors and used these gross-measurement-adjusted values for the flux balance analyses (**Section 1.13 and Data S4 [Simulations 25-28, 37-40, and 49-52], ESI†**). The GEMs found feasible solutions with a 0.1%, 0.1%, and 0.5% deviation of the reaction flux bounds from the calculated fluxes for *M. thermautotrophicus* ΔH, *M. thermautotrophicus* Z-245, and *M. marburgensis* Marburg, respectively (**Data S4, ESI†**). The maximum fluxes of the non-growth-associated maintenance energy (reaction ATPM) varied between the three microbes. The non-growth-associated ATP maintenance energy represents the dissipation of ATP (as no storage compounds are known).^58^ It is highest for *M. thermautotrophicus* ΔH, compared to *M. thermautotrophicus* Z-245 and *M. marburgensis* Marburg (**Table 4**, with the assumption that the biomass compositions are the same for the three microbes). The higher the non-growth-associated maintenance energy, the more methane (and ATP) must be produced per biomass unit. We further constrained the GEMs with transcriptomics and proteomics using the GIMME algorithm,^59^ which resulted in reduced GEMs. We performed a set of flux balance analysis simulations comparing the unconstrained and reduced GEMs (**Figure 3**; **Data S4 [transcriptomics: Simulations 29-32, 41-44, and 53-56; proteomics: Simulations 33-36, 37-40, and 57-60] and Section 1.25, ESI†**). This allowed us to have a close look at specific pathways. For example, we looked at the conversion of methenyl-H_4_MPT to methylene-H_4_MPT, but this did not resolve the exact flux catalyzed by different enzymes for these reactions (**Text S3, ESI†**). However, we were able to detect pathway differences in the formate anabolism between the three microbes.

**Table 4.** ATPM flux ranges for different constraints (ranging from biomass production maximization to ATPM maximization).

| DH<sup>1</sup> | ZZ<sup>2</sup> | MM<sup>3</sup> |
| --- | --- | --- |
| Model<sup>4</sup>: 1.5-17.85 | Model: 1.5-10.73 | Model: 1.5-12.20 |
| Model trans<sup>5</sup>: 1.5-17.46 | Model trans: 1.5-10.48 | Model trans: 11.96-12.19 |
| Model prot<sup>6</sup>: 1.5-17.55 | Model prot: 1.5-10.44 | Model prot: 1.5-11.90 |

815 <sup>1</sup> DH, *M. thermotrophicus* ΔH816 <sup>2</sup> ZZ, *M. thermotrophicus* Z-245817 <sup>3</sup> MM, *M. marburgensis* Marburg818 <sup>4</sup> Model, model only constrained with experimental data819 <sup>5</sup> Model trans, model additionally constraint with the transcriptomics data820 <sup>6</sup> Model prot, model additionally constraint with the proteomics data

In all three microbes, formate is a crucial precursor for purine biosynthesis and has to be synthesized during growth with molecular hydrogen and carbon dioxide.^60–62^ With our pan-genome comparison, we found the formate acetyltransferase/pyruvate-formate lyase (Pfl) (PFL reaction) in *M. thermautotrophicus* ΔH (MTH346/ISG35_1600), which was absent in both *M. marburgensis* Marburg^19,63^ and *M. thermautotrophicus* Z-245. This enzyme catalyzes the CoA-dependent reversible conversion of pyruvate to formate (for modeling purposes, the reaction is used irreversibly due to thermodynamics considerations, **Section 1.6, ESI†**). We found several putative Pfl-activating enzymes in the three microbes (**Text S4, ESI†**). However, given the missing Pfl, we assumed that *M. thermautotrophicus* Z-245 and *M. marburgensis* Marburg do not utilize the PFL reaction. Thus, no genes are included for those reactions in the GEMs.

Theoretically, in addition to the Pfl, *M. thermautotrophicus* ΔH can use a formate dehydrogenase (Fdh, ISG35_05415 and ISG35_05410, gene groups 967 and 968) to synthesize formate for biomass growth. The Fdh was previously hypothesized to be the relevant reaction for formate production.^19^ However, reducing the GEM with proteomics data resulted in the Pfl as the preferred formate production route because Fdh was not detected with proteomics, thus, eliminating the corresponding FDHF420 reaction in the flux balance analysis (**Figure 3; Data S4, ESI†**). The Pfl level was also elevated compared to the Fdh in the transcriptomics data (**Figure 3; Data S1, ESI†**). This implied that *M. thermautotrophicus* ΔH primarily produced formate for biomass growth *via* Pfl. This hypothesis will need to be confirmed experimentally to corroborate the proteomics-derived evidence for the lack of the Fdh enzyme.

In contrast to both *M. thermautotrophicus* ΔH and *M. marburgensis* Marburg, *M. thermautotrophicus* Z-245 has two Fdhs. The first Fdh is encoded in the operon *fdhCAB* (i.e., *fdh_cassette_*, ISG36_07620, ISG36_07615, ISG36_07610), which includes a putative formate transporter (FdhC). This Fdh was previously reported to be responsible for the utilization of formate as an alternative carbon and electron source.^21^ Nölling and Reeve^21^ found upregulation of expression of the *fdh_cassette_* when *M. thermautotrophicus* Z-245 cells were limited by molecular hydrogen availability. Interestingly, despite a high molecular hydrogen availability in our bioreactors, we obtained a high absolute abundance of the Fdh_cassette_ in our transcriptome (9.25 ± 1.52, 33.35 ± 2.44, and 0.26 ± 0.08 transcripts per million for *fdhA, fdhB*, and *fdhC*, respectively) and proteome (1.16E+08 ± 1.26E+07, 1.55E+08 ± 1.34E+07, and 7.66E+07 ± 2.19E+07 iBAQ for FdhA, FdhB, and FdhC, respectively) (**Figure 3**; **Data S1, ESI†**). This indicated that the Fdh_cassette_ might catalyze the reverse reaction during growth on molecular hydrogen and carbon dioxide, as was discussed by Nölling and Reeve^21^.

The second Fdh in *M. thermautotrophicus* Z-245 is encoded by the gene groups 967 and 968, and there is no putative transporter-encoding gene in the genetic vicinity. This second Fdh is homologous to the Fdh found in *M. thermautotrophicus* ΔH and *M. marburgensis* Marburg. In *M. thermautotrophicus* Z-245, the transcript levels of this second Fdh were less abundant (1.13 ± 0.31 vs. 9.25 ± 1.52 [fdhA] and 11.88 ± 2.68 vs. 33.35 ± 2.44 [fdhB] TPM) compared to the Fdh_cassette_ and it was not detected with proteomics. This indicated that in *M. thermautotrophicus* Z-245, the Fdh_cassette_ might also be responsible for formate production for biomass growth. Indeed, our flux balance analysis simulations with *M. thermautotrophicus* Z-245 primarily predicted the use of the FDH_F420 (Fdh_*cassette*_) in the direction of formate production rather than FDHF420 (second Fdh; including the omics-constrained runs, except for the loopless flux balance analysis with transcriptomics) (**Figure 3**; **Data S4, ESI†**).

Because *M. marburgensis* Marburg neither encodes the Pfl nor the Fdh_cassette_, formate is likely only produced from the second Fdh of gene groups 967 and 968. This was also hypothesized previously, and this hypothesis was supported by an auxotrophic formate strain of *M. marburgensis* Marburg, which did not exhibit Fdh activity anymore.^64^ All flux balance analysis simulations (including the omics-constrained ones) predicted using the FDHF_420_ reaction (**Data S4, ESI†**). The two subunits were upregulated in the transcriptome by 2.4-4.9 log2FC compared to the other two microbes (**Figure 3; Data S1, ESI†**). However, we found only one subunit (FdhA) in the proteome with low abundance (gene group 967: 0.000132%). This was three orders of magnitude lower than the homologous subunit of the Fdh_cassette_ in *M. thermautotrophicus* Z-245 (FdhB: 0.254%, FdhA: 0.191%, FdhC: 0.126%). Theoretically, the proteome should better depict the reality that is occurring in the cell than the transcriptome. However, the proteome is more challenging to measure and analyze than the transcriptome.^54^ Because *M. marburgensis* Marburg reached higher biomass production rates (**Figure 1C**), these findings raise questions on how *M. marburgensis* Marburg produced formate for biomass growth. Nevertheless, our analysis may indicate that the three methanogens have each evolved to prefer a different pathway to produce the necessary formate for biomass growth.

Potentially, formate production for *M. marburgensis* Marburg was achieved by other yet unconsidered means. If the low-abundant Fdh was responsible for formate production, questions on the kinetic properties of this Fdh remain. Alternatively, a hydrogen-dependent carbon dioxide reductase (HDCR)-like activity might provide the formate for anabolism.^65^ While formate is not an intermediate in carbon dioxide reduction during methanogenesis, a weak formate dehydrogenase activity has been found for Fmd/Fwd.^66^ Recently, a large enzyme complex was identified to be involved in the electron-bifurcating step of carbon dioxide reduction without the release of ferredoxin.^67^ This enzyme complex might provide the formate that is required for anabolism by an apparent HDCR-like activity. While more experimentation will be required to resolve the details of formate anabolism in *M. marburgensis* Marburg, we still investigated the change in the (formate) metabolism and the role of the Fdh_cassette_ in a genetically engineered *M. thermautotrophicus* ΔH strain that encodes the Fdh_cassette_ from *M. thermautotrophicus* Z-245 on a plasmid (*M. thermautotrophicus* ΔH pMVS1111A:P_*hmtB*_-*fdh*_Z-245_).^40^

### Genetic engineering allows stable growth of *M. thermautotrophicus* ΔH on formate

We compared the ability of *M. thermautotrophicus* ΔH pMVS1111A:P_*hmtB*_-*fdh*_Z-245_ to grow on formate^40^ to the native formate-utilizer, *M. thermautotrophicus* Z-245. We operated chemostat bioreactors that were fed with 355 ± 5 mM sodium formate (**Figure 4A**; **Section 1.8, ESI†**). We again excluded cross-contamination between the bioreactors with strain-specific PCR analyses (**Figure S6 and ESI† Section 1.9, ESI†**). The two microbes did not have significantly different sodium formate uptake rates or methane, carbon dioxide, and biomass production rates (**Figure 4B; Data S5, ESI†**). However, the molecular hydrogen production rate of *M. thermautotrophicus* ΔH pMVS1111A:P_*hmtB*_-*fdh*_Z-245_ was significantly higher (1.53-fold) than that of *M. thermautotrophicus* Z-245, albeit both were low (**Figure 4**). Thus, the gas data indicated no significant disadvantage to harboring the plasmid for growth on sodium formate. Notably, the plasmid was maintained in the culture without adding antibiotics (**Figure S6, ESI†**). Also for *M. thermautotrophicus* ΔH pMVS1111A:P_*hmtB*_-*fdh*_Z-245_ that was grown on molecular hydrogen and carbon dioxide without antibiotics, the plasmid was found in the bioreactor at the end of the experiment. While we cannot completely rule out that only a subpopulation maintained the plasmid, this finding further demonstrates the applicability of the genetic system for metabolic engineering purposes. The slightly lower uptake rates and the higher biomass production rates of the genetically modified strain were more similar to those of *M. thermautotrophicus* Z-245 and *M. marburgensis* Marburg than to wild-type *M. thermautotrophicus* ΔH (**Figure S7, ESI†**). This shows that, indeed, the different solutions for each of the three methanogens to produce formate in anabolism results in different methane-to-biomass ratios, which is an important parameter in bioprocessing (it should be high). Of course, further work is necessary to validate this hypothesis. We collected additional omics data and, while out of the scope of this study, will analyze these data in combination with our GEMs to investigate potential impacts of the plasmid and episomal formate dehydrogenase on rearrangements in the metabolism of *M. thermautotrophicus* ΔH pMVS1111A:P_*hmtB*_-*fdh*_Z-245_, also concerning formate anabolism.

**Figure 4.**
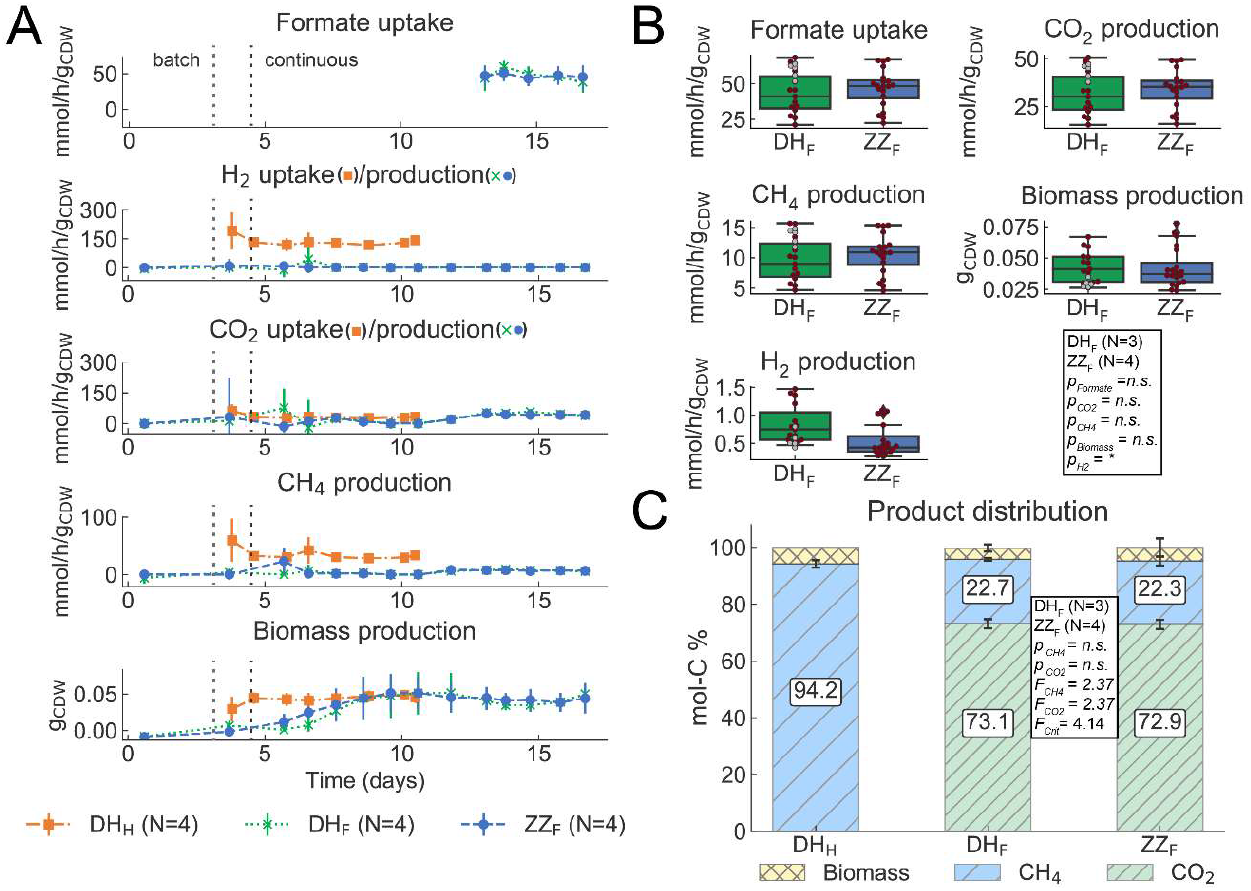
Second experiment fermentation data from chemostat bioreactors with *M. thermautotrophicus* ΔH pMVS1111A:P_*hmtB*_-*fdh*_Z-245_ (DH) and *M. thermautotrophicus* Z-245 (ZZ). **A)** Gas consumption (molecular hydrogen and carbon dioxide uptake) or sodium formate consumption, and methane and biomass production data from quadruplicate bioreactors for the fermentation period of 11 (DH_H_) and 17 (DH_F_ and ZZ_F_) days. Formate uptake rates were only determined once steady-state was reached and all formate provided was consumed. **B)** Average sodium formate consumption and molecular hydrogen, carbon dioxide, methane, and biomass production data during the steady-state period (days 13 to 17) for DH_F_ and ZZ_F_. For statistical analysis in pair-wise comparisons with t-test, data points without suspected gross measurement error (red circles) were included, and data points with suspected gross measurement error (gray circles) were excluded (**Section 1.13, ESI†**). **C)** Average normalized product distribution, including a statistical analysis by ANOVA for DH_F_ and ZZ_F_ (**Section 1.14, ESI†**). CH_4_, methane; CO_2_, carbon dioxide; H_2_, molecular hydrogen; DH, *M. thermautotrophicus* ΔH pMVS1111A:P_*hmtB*_-*fdh*_*Z*-245_; ZZ, *M. thermautotrophicus* Z-245; F, sodium formate as substrate; H, molecular hydrogen and carbon dioxide as substrates; *, *p* < 0.05; n.s., not significant (p > 0.05); *F, F* value; F_crit_, *F* critical value.

## Summary and Outlook

Power-to-gas provides a means for storing excess renewable electric power while simultaneously capturing carbon dioxide. Methanogens can act as biocatalysts to generate methane. Although methanogenesis is well studied, there are still knowledge gaps surrounding methanogenic metabolism, which limit their ability to be harnessed for more versatile biotechnological applications.

In this study, we compared *M. thermautotrophicus* ΔH, *M. thermautotrophicus* Z-245, and *M. marburgensis* Marburg at a genomic, transcriptomic, and proteomic level to identify differences in their metabolism. While the genomes encode more than 1600 homologs, most genes were differentially expressed at the transcriptomic level (**Table 3**). In contrast, the proteome did not exhibit nearly as many differences. We note that in both omics datasets, fewer differences were found between the two *M. thermautotrophicus* species compared to *M. marburgensis* Marburg, which is consistent with their phylogeny.^18^ Under the conditions of our bioreactor experiments, *M. thermautotrophicus* ΔH had a higher specific methane production rate than the other microbes, while *M. marburgensis* Marburg reached higher biomass production rates. From our modeling results (with the maximization of the ATPM reaction), it was possible to deduce that *M. thermautotrophicus* ΔH has the highest and *M. marburgensis* Marburg has the lowest non-growth-associated maintenance energy. As ATP is the primary energy currency in the cell, this is an important finding to facilitate adopting and optimizing *Methanothermobacter* spp. as cell factories for different purposes.^68^ While we could not ascertain a clear reason for this difference, our multi-omics analysis led us to propose that differences in anabolism could explain this observation. We identified that the three microbes might use three different enzymes to produce formate for biomass growth, such as for purine biosynthesis.^60–62^ Potentially, these differences in formate anabolism can have an impact on the adaptation to different ecological niches in which, for example, a higher methane production rate (faster kinetics) over biomass yields would provide a selective benefit for *M. thermautotrophicus* ΔH. Further investigations are required to test this hypothesis.

Understanding the metabolism of a microbe is especially important to guide its rewiring for biotechnological purposes (*e.g*., the use of different substrates or the production of value-added compounds). Our recently developed genetic system for *M. thermautotrophicus* ΔH demonstrated gain-of-function modifications with a shuttle-vector system.^40^ We had shown that adding the Fdh_cassette_ from *M. thermautotrophicus* Z-245 to *M. thermautotrophicus* ΔH permitted growth on formate as the sole carbon and electron source (natively, *M. thermautotrophicus* ΔH only utilizes carbon dioxide and molecular hydrogen).^40^ Here, we demonstrated the long-term ability of the resulting strain *M. thermautotrophicus* ΔH pMVS1111A: P_*hmtB*_-*fdh*_Z-245_ to grow on formate and achieve a performance that is comparable to the performance of wild-type *M. thermautotrophicus* Z-245 without antibiotic selection. The ability to combine the GEM with the genetic system will be a considerable step toward chemical production in power-to-x applications (compared to power-to-gas, power-to-x has a more flexible product spectrum that is not limited to only molecular hydrogen or methane^69^) with thermophilic methanogens because now the verified and validated GEM can be applied to predict phenotypes.

*Methanothermobacter* spp. have already been implemented for power-to-gas applications. Understanding the metabolic strengths and limitations of different strains will help to select the most appropriate strain for a given application. Here, we found that *M. thermautotrophicus* ΔH has the highest methane-to-biomass ratio, which is beneficial for power-to-gas applications. Applying our genetic system, we demonstrate that amending the metabolism of *M. thermautotrophicus* ΔH with a catabolic formate dehydrogenase changes the mode of formate anabolism, which supports our hypothesis of different enzymes for formate anabolism by the three microbes. This finding has implications to formulate strategies for the redirection of carbon from methane (and biomass) to other high-value products with metabolic engineering strategies.

## Data availability

The **genome** sequences, annotations, and methylation patterns were deposited to NCBI ^70^ and can be found under Bioproject ID PRJNA674001 (**Table S1**). The **transcriptomics** data (gene expression data) discussed in this publication have been deposited in NCBI’s Gene Expression Omnibus^71^ and are accessible through GEO Series accession number GSE218145 (https://www.ncbi.nlm.nih.gov/geo/query/acc.cgi?acc=GSE218145). The **proteomics** DDA data are in the submission process to be deposited to the ProteomeXchange Consortium *via* the PRIDE partner repository.^72–74^ The **modeling**-related files (**Data S2, ESI†**) were deposited in an Open Modeling EXchange format (OMEX)^75^ to BioModels^76,77^ and assigned the identifiers MODEL2211290001 (M. *thermautotrophicus* ΔH), MODEL2211290002 (M.

*thermautotrophicus* Z-245), and MODEL2211290003 (M. *marburgensis* Marburg). The pan-model is provided as an Excel^®^ model (.xlsx, densely commented). The GEMs for each microbe are provided in the following formats: .xml (SBML^78^), .json, and .mat. Additional files include: the benchmarking reports (.html) from MEMOTE,^46^ the FROG analysis files (.omex) from fbc_curation,^79^ and the map files for visualization (.json) with Escher.^48^ Further, the data used to constrain the models is provided as supplementary data contained in **Data_SX.zip, ESI†.** This includes: the gas fermentation data (as Excel^®^ files, **Data S3** and **Data S5, ESI†**) and the individual microbe and differential expression analyses data for both the transcriptomics and proteomics (as Excel^®^ files, **Data S1, ESI†**). Additional scripts and programs (**Data S6, ESI†**) are available on GitHub at https://github.com/isacasini/Casini_2022_GEM.

## Acknowledgments

This work was supported by the Humboldt Foundation in the framework of the Humboldt professorship, which was awarded to LTA, and by the Max Planck Institute for Biology Tübingen (REL and LTA). IC would like to acknowledge support from the German Academic Exchange Service (DAAD) through the DAAD *Kurzzeitstipendien für Doktoranden*. EM acknowledges support from the Australian Research Council for the Centre of Excellence in Synthetic Biology. The authors would like to thank Luis Antoniotti and Jürgen Barth from the Max Planck Institute for Biology Tübingen workshop for their help with modifying the bioreactor system. The authors would like to thank Lucas Mühling, Nils Rohbohm, and Andrés Ortíz Ardilla for their helpful input and support with experimentation and Nicolai Kreitli for support with bioreactor experiments. The authors would like to acknowledge the assistance of the Genome Center of the Max Planck Institute for Biology Tübingen, particularly Christa Lanz, for their assistance in the PacBio sequencing, and the Proteome Center Tübingen (PCT) for their support with the proteomics. The authors would like to acknowledge the fruitful conversations with Theresa Ahrens and Doris Hafenbradl from Electrochaea GmbH, Planegg, Germany. The graphical abstract was created with Biorender.

## CRediT Author Contributions

**Isabella Casini:** Methodology, Software, Formal Analysis, Investigation, Data Curation, Writing - Original Draft & Review & Editing, Visualization, Project Administration, Funding acquisition; **Tim McCubbin:** Methodology, Software, Formal Analysis, Writing - Review & Editing; **Sofia Esquivel-Elizondo:** Methodology, Investigation, Writing - Review & Editing; **Guillermo G. Luque:** Software, Formal Analysis, Writing - Review & Editing; **Daria Evseeva:** Formal Analysis, Investigation, Writing - Review & Editing; **Christian Fink:** Investigation, Writing - Review & Editing; **Sebastian Beblawy:** Investigation, Writing - Review & Editing; **Nicholas D. Youngblut:** Methodology, Writing - Review & Editing; **Ludmilla Aristilde:** Writing - Review & Editing; **Daniel H. Huson:** Supervision, Writing - Review & Editing, Funding Acquisition; **Andreas Dräger:** Software, Writing - Review & Editing, Funding Acquisition; **Ruth E. Ley:** Resources, Supervision, Writing - Review & Editing, Funding Acquisition; **Esteban Marcellin:** Methodology, Resources, Writing - Review & Editing, Funding Acquisition; **Largus T. Angenent:** Conceptualization, Resources, Writing - Review & Editing, Supervision, Project administration, Funding Acquisition; **Bastian Molitor:** Conceptualization, Writing - Original Draft & Review & Editing, Supervision, Project administration, Funding acquisition

