## Supplementary information for "An integrated systems-biology platform for power-to-gas technology"

### Electronic Supplementary Information

#### Table of Contents


### 1. Materials and Methods

1.1. Microbial strains and medium composition
1.2. Genome sequencing
1.3. Genome assembly
1.4. Genome comparisons
1.5. Clusters of orthologous genes (COG) functional annotation
1.6. Genome-scale metabolic model reconstruction
1.7. Biomass composition determination and maintenance energies
1.8. Bioreactor setup and operating conditions
1.9. Cross-contamination check
1.10. Fermentation gas analysis
1.11. Biomass concentration analysis
1.12. Sodium formate concentration measurements
1.13. Carbon and electron balance calculations
1.14. Normalized product distribution
1.15. Interspecies comparison
1.16. RNA sample preparation
1.17. RNA sequencing
1.18. Raw RNA sequencing data analysis
1.19. Protein sample preparation
1.20. Protein measurement
1.21. Protein analysis with proteome discoverer
1.22. Pan-genome differential expression database creation
1.23. Eha/Ehb ratio determination
1.24. Methanogenesis relative abundances
1.25. Integrating fermentation data in the GEMs

### 2. Results

2.1. Supplementary Text S1 to S4
2.2. Supplementary Figures S1 to S8

**2.3. Supplementary Tables Legends S1 to S15 (for Tables in tables.xlsx, ESI†)**

**2.4. Data Files Descriptions S1-S6 (including for Data in Data_SX.zip, ESI†)**

#### 3. Supplementary References

### 1. Materials and Methods

#### 1.1 Microbial strains and medium composition

*M. thermautotrophicus* ΔH (DSM 1053), *M. thermautotrophicus* Z-245 (DSM 3720), and *M. marburgensis* Marburg (DSM 2133) were obtained from the DSMZ (Braunschweig, Germany) and were essentially cultivated as described in Fink*, et al.*[1] For anaerobic handling of media and materials the atmosphere of the anaerobic chamber (UniLab Pro Eco, MBraun, Garching, Germany) contained 95% v/v $N_2$ and 5% v/v $CO_2$. *M. thermautotrophicus* ΔH pMVS1111A:$P_{hmtB}$-$fdh_{Z\text{-}245}$ was previously generated in our lab.[1] Preparation of batch and continuous media for bioreactor runs was adjusted from the mineral media of Balch*, et al.*[2] and Martin*, et al.*[3] The mineral medium contained (per liter): nitrilotriacetic acid (NTA), 0.096 g; trisodium nitrilotriacetate, 0.275 g; sodium chloride, 0.675 g; di-potassium hydrogen phosphate, 0.255 g; ammonium chloride, 2.006 g; magnesium chloride hexahydrate, 0.12 g; calcium chloride dihydrate, 0.090 g; potassium di-hydrogen phosphate, 0.345; ammonium nickel sulfate, 1.5 mL (0.2% w/v); iron(II) chloride tetrahydrate, 1.5 mL (0.2% w/v); resazurin indicator solution, 4 mL (0.025% w/v); and trace element solution, 1.5 mL. The trace element solution was prepared 10-fold as stated by Balch*, et al.*[2] with minor modifications, and contained (per liter): NTA, 2.0 g; magnesium sulfate heptahydrate, 30.0 g; manganese(II) sulfate, 5.0 g; sodium chloride, 10.0 g; iron(II) sulfate heptahydrate, 1.0 g; cobalt(II) chloride hexahydrate, 1.8 g; calcium chloride dihydrate, 1.0 g; zinc sulfate heptahydrate, 1.8 g; copper sulfate pentahydrate, 0.1 g; aluminum potassium sulfate dodecahydrate, 0.18 g; boric acid, 0.1 g; sodium molybdate dihydrate, 0.1 g; ammonium nickel(II) sulfate hexahydrate, 2.8 g; sodium tungstate dihydrate, 0.1; and sodium selenate, 0.1 g. The pH of the trace element solution was adjusted to 6.0 with 5 M potassium hydroxide. In all the continuous media, 0.02 mL/L of Anti Foam SE-15 (Sigma-Aldrich, Merck, Darmstadt, Germany) was supplemented. For growth with sodium formate, serum bottles for batch cultivation were sparged with $N_2/CO_2$ (80/20 %, v/v), and 100 mM sodium formate was added after autoclaving. Further, the serum bottles were supplemented with 0.001 mM sodium selenate and 0.01 mM sodium molybdate dihydrate. For growth with sodium formate, the continuous media contained 355 ± 5 mM sodium formate and was supplemented with 0.0015 mM sodium selenate and 0.015 mM sodium molybdate dihydrate (final pH was 5.85). A concentrated sodium formate solution was prepared and sterilely added to the continuous media after autoclaving, sparging, and reducing the media. All media were prepared with Millipore water (18.2 MΩ·cm). The medium was autoclaved either in Schott bottles with butyl septa with Masterflex® L/S Norprene Food-Grade Tubing, L/S 14 tubing (Cole-Parmer GmbH, Wertheim, Germany) or directly in the bioreactor vessels.

#### 1.2 Genome sequencing

*M. thermautotrophicus* ΔH, *M. thermautotrophicus* Z-245, and *M. marburgensis* Marburg were grown in mineral medium overnight. The entire biomass (50 mL) was collected by centrifugation at 3170 × g and room temperature for 20 min (5920 R Eppendorf, Hamburg, Germany), and the genomic DNA was extracted using a phenol-chloroform extraction method. For this, to the biomass pellet, 500 µL of cetyltrimethylammonium bromide buffer (CTAB, made according to Cold Spring Harbor Protocols[4], however, without the Polyvinylpyrrolidone) was added, and the pellet was resuspended. The mixture was transferred to sterile 2 mL bead beating tubes (containing 500 µL of 0.1 µm BeadBeater® zirconia beads, Carl Roth, Karlsruhe, Germany) and vortexed (Vortex-Genie® 2, VWR International GmbH, Darmstadt, Germany) at 2700 $min^{-1}$ for 5 s on, then 1 s off, repeatedly for 1 min, after which the tube was placed on ice. 500 µL of ROTI®Phenol/Chloroform/Isoamyl alcohol (ratio of 25:24:1, Carl Roth, Karlsruhe, Germany) was added, the tube inverted, and then centrifuged at 4°C and 16000 × g for 10 min (5424, Eppendorf, Hamburg, Germany). The top layer was transferred (using wide orifice pipette tips) to a phase lock tube, which was prepared ahead by adding 2 $mm^3$ of vacuum grease Dow Corning® (VWR International GmbH, Darmstadt, Germany) into 2 mL tubes, which were centrifuged until 9391 × g was reached, and then autoclaved to sterilize. Another 500 µL of ROTI®Phenol/Chloroform/Isoamyl alcohol was added, the tube was inverted to mix, and centrifugation was performed as in the previous step. The supernatant was transferred to new tubes, 500 µL of Chloroform/Isoamyl alcohol (24:1, VWR International GmbH, Darmstadt, Germany) was added, the tube was inverted to mix, and centrifugation was performed as in the previous step. The supernatant was added to a fresh tube, and the gDNA was precipitated by adding 0.1 volumes of cold 3 M sodium acetate and 2 volumes of ice-cold absolute ethanol, inverting the tube to mix, and then incubating it overnight at -20°C. The following day, the tube was centrifuged again at 4°C and 16000 × g for 10 min, the supernatant was removed, and the gDNA was washed with 300 µL of ice-cold 70% v/v ethanol. Centrifugation was performed as in the previous step, the supernatant was removed, and the tube was air-dried for approximately 1.5 h at 50°C in a ThermoMixer® C (Eppendorf, Hamburg, Germany). The pellet was resuspended with 44 µL elution buffer (10 mM Tris·Cl, pH 8.5, and nuclease-free water [New England Biolabs, Ipswich, United States]) and allowed to rest for an hour to resolve at room temperature. To remove RNA, 1 µL of Bovine Ribonuclease A (VWR International GmbH, Darmstadt, Germany) was added, and the tube was allowed to rest for 30 min at room temperature. The quality of the gDNA was then checked using the Femto Pulse System (1.0.0.32, Agilent, Santa Clara, United States) according to the manufacturer's instructions, using the Genomic DNA 165 kb Kit (Agilent, Santa Clara, United States) and a 70 min separation time. The quantity of double-stranded DNA was measured with the Qubit® 2.0 Fluorometer (Invitrogen, Carlsbad, CA, USA) using the Qubit® dsDNA HS Assay Kit (Thermo Fischer Scientific, Dreieich, Germany). Library preparation was conducted with the SMRTbell® Express Template Preparation Kit (PN 101-397-100 Version 3, January 2018, Pacific Biosciences, Menlo Park, United States) as specified in the manufacturer's instructions. The genomes were then sequenced using the Sequel I System (Pacific Biosciences, Menlo Park, United States).

##### 1.3 Genome assembly

*De-novo* genome assembly (including the plasmid when applicable) was conducted, and methylation patterns were deciphered from the PacBio sequencing results. The utilized pipeline included seven main aspects: **1)** contamination control by DIAMOND alignment against the NCBI-nr database to confirm the absence of any non-*Methanothermobacter* reads; **2)** assembly by Canu (which includes error correction, trimming, and assembly); **3)** first polishing by BlastR, samtools, and Arrow; **4)** circularizing by Blast and BioPython; **5)** second polishing by BlastR, samtools, and Arrow; and **6)** methylation patterns prediction by ipdSummary, MotifMaker, and BaseModFunctions. The final annotation was performed by National Center for Biotechnology Information (NCBI) upon genome sequence submission. Details on the applied programs can be found in **Table S12, ESI†**.

#### 1.4 Genome comparisons

Protein FASTA files were generated using the GenBank files from NCBI and the BioPython version 1.77.[5] Genome-wide comparisons were performed with protein Basic Local Alignment Search Tool (BLASTp+) version 2.10.0 locally.[6-8] The new *M. thermautotrophicus* ΔH and *M. marburgensis* Marburg sequences were compared to the previously published ones, accession numbers `NC_000916.1`[9] and `NC_014408.1`[10], respectively. The alignments were performed twice, both times with an expectation value (*e*-value) cut-off of 10E-30. The first time, the new sequences were set as the queries and the old sequences as the subjects. The second time the query and subject were reversed. The protein hit with the highest bit score was taken as the matching protein (and gene). For homologous gene comparisons (which were used for the pan-genome and pan-model construction), the *e*-value cut-off was set to 0.001. The following pairwise comparisons were performed with the following query and subject pairs: **1)** *M. thermautotrophicus* ΔH and *M. marburgensis* Marburg; **2)** *M. thermautotrophicus* ΔH and *M. thermautotrophicus* Z-245; and **3)** *M. marburgensis* Marburg and *M. thermautotrophicus* Z-245. The protein hit with the highest bit score was taken as the homologous protein (and gene). The three comparisons were merged with *M. thermautotrophicus* ΔH as the backbone *via* the `LOOKUP` function in Excel®. Some supplementary gene-wise annotations were completed with the online BLASTp tool.[11]

#### 1.5 Clusters of orthologous genes (COG) functional annotation

The COG functional annotation was performed on the protein FASTA file for each microbe with the script `cdd2cog` v0.2 from the bac-genomics-scripts according to the author's instructions.[12] A modification to the code, as suggested in issue #14 (https://github.com/aleimba/bac-genomics-scripts/issues/14, accessed on 12/2021), was applied to allow the use of the COG2020 database.[13]

#### 1.6 Genome-scale metabolic model reconstruction

A pan-model for *M. thermautotrophicus* ΔH, *M. thermautotrophicus* Z-245, and *M. marburgensis* Marburg, which included all reactions from the three microbes, was built in Microsoft® Excel® (Microsoft 365 MSO, Version 2202, Washington, United States), following the protocol established by Thiele and Palsson[14]. The pan-model used genome sequences, assemblies, and annotations as its backbone. Reactions and pathways were added based on data from KEGG,[15] ModelSEED,[16] UniProt,[17] Brenda,[18] BioCyc,[19] MetaCyc,[20] BIGG,[21] and NCBI.[22] Metabolite protonation was determined (for pH 7.0) using the command line tool `cxcalc` and MarvinSketch 18.8.0, ChemAxon (http://www.chemaxon.com, accessed in 2018). When possible, reactions and genes were verified with literature. Genes for which no genus-specific evidence was found were BLAST searched to genes of species with stronger evidence. These BLAST results are specified in the comment section of the reconstruction. Over 790 references were cited for which the microbe and type of evidence were recorded using Evidence & Conclusion Ontology (categories: biochemical, genetic, physiological, sequence, modeling, and no data), which were then used to determine the confidence of each reaction for each microbe.[14] Published GEMs from the following microbes were used to gap-fill and validate pathways in the *Methanothermobacter* pan-model: **1)** *Methanosarcina acetivorans*: *i*VS941,[23] *i*MB745,[24] *i*MAC868a,[25] *i*ST807;[26] **2)** *Methanosarcina barkeri*: *i*AF692,[27] *i*MG746;[28] **3)** *Methanospirillum hungatei*: *i*Mhu428;[29] **4)** *Methanococcus maripaludis* S2: *i*MM518;[30,31] **5)** *Methanobrevibacter smithii*: *i*Msi385;[32] **6)** *Methanocaldococcus jannaschii*: *i*TS436.[33] The pan-model was converted to three strain-specific GEMs written in SBML Level 3 Version 1,[34] including the extension packages for flux balance constraints (fbc) version 2[35] and groups,[36] and verified in MEMOTE 0.13.0.[37] For verification in MEMOTE, the GEMs were constrained with Conditions 1, 8, and 15 for *M. thermautotrophicus* ΔH, *M. thermautotrophicus* Z-245, and *M. marburgensis* Marburg, respectively (**Data S2 and Data S4, ESI†**). The GEMs aimed to be MIRIAM-compliant, including metabolite and reaction annotations with Compact Identifiers[38] for various databases (gene annotations were not available given the newly annotated genomes. However, the old gene annotations can be found in **Data S2, ESI†**). The directionality of the reactions was determined using thermodynamics-based flux variability analysis (TFVA)[39] and the ModelSEED database.[40] For the TFVA method, a sink reaction (reversible uptake and production possible) was added to each metabolite in the GEMs. A core model was then generated using the `tmodel` function with the following constraints for cytosol and extracellular: **1)** pH (7.6, 7.3); **2)** ionic strength (0.1, 0); **3)** temperature (338.15 K, 338.15 K); and **4)** membrane potentials ([0,150], [-150, 0] for [cytosol, extracellular]). The minimum and maximum flux values were found with Gurobi as the solver, using the box or univariate method.[39] If the flux values spanned zero, the reaction bounds were set to (-1000, 1000). If both the minimum and maximum flux values were less than zero, the reaction bounds were set to (-1000, 0). Lastly, if both the minimum and maximum flux values were greater than zero, the reaction bounds were set to (0, 1000). If directionally could not be determined with either TFVA or the ModelSEED database, directionality was set to reversible, except for reactions that caused loops with ATP (*e.g.*, with ATP losing a phosphate group,[14] or between redox carriers[41]). Seven constrain conditions per microbe were designed that depict relevant cultivation conditions: Conditions 1-7, Conditions 8-14, and Conditions 15-21 for *M. thermautotrophicus* ΔH, *M. thermautotrophicus* Z-245, and *M. marburgensis* Marburg, respectively (**Data S4, ESI†**). Flux balance analyses (FBAs) with maximization of biomass exchange (`EX_biomass_e`) as the objective function and these different conditions were performed to verify the ability of the three strain-specific GEMs to grow under these cultivation conditions (**Data S4**

**[Simulations 1-24], ESI†)**. The FBAs were run using COBRApy version 0.22.1 (**Data S4 and Data S6, ESI†**).[42]

#### 1.7 Biomass composition determination and maintenance energies

The biomass composition for the three microbes in the GEMs is assumed to be the same (**Data S2, ESI†**). Briefly, the fraction for each of the molecules that make up the biomass was found by averaging those taken from previously published methanogen GEMs[43] and the incorporation of empirically found data for *M. thermautotrophicus* ΔH (**reaction ID, `BIOMASS_X`, where X is DH, ZZ, or MM; Data S2, ESI†**). The elemental composition of the biomass, $CH_{1.681}O_{0.418}N_{0.222}S_{0.004}$ (molecular weight of 23.502 g/mol) was taken from Duboc*, et al.*[44] Growth-associated maintenance (GAM) energy is included in the biomass macromolecule synthesis reactions. Non-growth-associated ATP maintenance costs (NGAM) are represented using the ATP hydrolysis reaction (`ATPM`).

#### 1.8 Bioreactor setup and operating conditions

Continuous fermentations were carried out in the BioXplorer 100 bioreactor platform controlled with the WinISO version 2.3.149.1 software (H.E.L, London, England). Each bioreactor was equipped with temperature, pH (part number Z001013510), and ORP (part number Z061013510) sensors (I&L Biosystems GmbH, Königswinter, Germany); a 0.15 µm sparging stone; a magnetic coupled stirring system; three peristaltic pumps for media feed-in, base feed-in (1 N NaOH was used for pH control), and effluent-out; a mass-flow controller (Red-y smart min; Vögtlin, Muttenz, Switzerland) to control the inlet gas flow rate; a condenser for the exhaust-gas line; and a separate sampling and inoculum port (fitted with a rubber butyl stopper). The bioreactors were fitted with Masterflex® L/S Norprene Food-Grade Tubing, L/S 14 (Cole-Parmer GmbH, Wertheim-Mondfel, Germany) except for the gas inlet lines, which were fitted with Masterflex® C-Flex ULTRA tubing, L/S 16 (Cole Parmer, Wertheim-Mondfel, Germany). The upstream gas mixture was set using Bronkhorst EL-Flow® Prestige mass flow controllers (Bronkhorst Deutschland Nord GmbH, Kamen, Germany) and mixed in a doubledended cylinder (Swagelok® Stuttgart, Reutlingen, Germany). The exhaust gas flow rate was measured offline using a MilliGascounter MGC-1 V3.4 PMMA (Dr.-Ing. RITTER Apparatebau GmbH & Co. KG, Bochum, Germany). The pH and ORP sensors, the pumps, and the MFCs were calibrated before each experiment. The bioreactors were filled with the mineral medium and autoclaved for one hour at 121°C. Afterward, the bioreactors were connected to the bioreactor platform, and the temperature was set to 65°C with agitation at 700 rpm.

For the first bioreactor experiment, the bioreactors were then sparged through sterile filters Minisart® HY (0.2 µm pore size; Sartorius AG, Göttingen, Germany) for two hours with $H_2/CO_2$ (80/20% v/v) at a gas flow rate of 10 mL/min. Before inoculation, the mineral medium was reduced with sterile anaerobic L-cysteine-HCl (0.5 g/L) and disodium sulfide nonahydrate (0.3 g/L), and the pH control was set to 7.3. Each bioreactor was inoculated with either 4 mL (*M. thermautotrophicus* ΔH and *M. thermautotrophicus* Z-245) or 3.6 mL (*M. marburgensis* Marburg) of preculture grown in serum bottles to an $OD_{600}$ of 0.35-0.36. The bioreactors were operated in batch mode for one day until an $OD_{600}$ of approximately 1.00 was reached, at which point continuous mode was started with a medium feed at a dilution rate of 0.83 $d^{-1}$. The first continuous operating period was conducted for approximately 12 days (**Figure S8, ESI†**), after which the bioreactors were emptied except for about 3-5 mL, which were used as inoculum for a second period with a starting $OD_{600}$ of 0.20-0.30. When an $OD_{600}$ of approximately 1.00 was reached, the bioreactors were switched into continuous mode with a medium feed at a dilution rate of 1.11 $d^{-1}$. Steady-state was reached after three hydraulic retention times (HRT),[45] which was 2.7 days in our setup. After an additional 3 HRTs (day 6.8), the transcriptomics, proteomics, and gram cell-dry weight determination samples were taken.

For the second bioreactor experiment, the bioreactor setup was the same as in the first experiment, except that for bioreactors grown on sodium formate they were: **1)** pH adjusted using 1 N HCl instead of 1 N NaOH, and **2)** sparged with $N_2/CO_2$ (80/20 %, v/v) rather than $H_2/CO_2$ (80/20 %, v/v). Each bioreactor was inoculated with 6 mL (*M. thermautotrophicus* ΔH pMVS1111A:$P_{hmtB}$-$fdh_{Z\text{-}245}$ and *M. thermautotrophicus* Z-245) of preculture grown in serum bottles ($OD_{600}$ ~0.2). The bioreactors were operated in batch mode until an $OD_{600}$ of approximately 1.00 for bioreactors grown on molecular hydrogen and carbon dioxide and 0.10-0.15 for bioreactors grown on sodium formate was reached. At this point, the continuous mode was started with a media feed at a dilution rate of ~1.00 $d^{-1}$ (this rate was achieved by ramping over several days for the bioreactors grown on sodium formate). After maintaining steady-state for three HRTs (10.6 days for $H_2/CO_2$ and 16.8 days for sodium formate), samples for transcriptomics, proteomics, and gram cell-dry weight determination were taken.

Daily samples of $OD_{600}$, pH, exhaust gas flow rate, and inlet and exhaust gas composition were taken during both experiments and periods. For the liquid culture samples, 1 mL of dead volume was first removed, and then 1 mL was used for $OD_{600}$ and pH measurements. To measure pH, the samples were maintained at 65°C (ThermoMixer® 460-0223, Eppendorf, Hamburg, Germany) and measured within 1 min of taking the sample (FiveEasy™ Plus pH/mV Benchtop meter with the micro pH electrode LE422 [Mettler-Toledo GmbH, Gießen, Germany] calibrated at 25°C and set to 65°C).

#### 1.9 Cross-contamination check

The bioreactors were checked for cross-contamination by polymerase chain reaction (PCR) using the Thermo Scientific™ Phire Plant Direct PCR Master Mix (Thermo Fischer Scientific, Dreieich, Germany) and custom primers (**Table S13, ESI†**). Cells were lysed by boiling 300 µL of the bioreactor culture (with an approximate $OD_{600}$ of 1.00) at 100°C for 10 min (ThermoMixer® 460-0223, Eppendorf, Hamburg, Germany); 1 µL was directly taken as a template for the PCR reaction. Primers were used at a 10 µM concentration. The PCR was carried out for 28 cycles in a Mastercycler® pro S (Eppendorf, Hamburg, Germany). The results of the reactions were visualized with gel electrophoresis (1% w/v agarose and SYBR™ Safe DNA Gel Stain (Thermo Fischer Scientific, Dreieich, Germany) and Gel Doc™ XR+ visualizer (Bio-Rad, Feldkirchen, Germany).

#### 1.10 Fermentation gas analysis

A 490 Micro Gas Chromatograph (microGC; Agilent, Santa Clara, United States) fitted with a multi-valve port system (Teckso GmbH, Neukirchen-Vluyn, Germany) was used to analyze the inlet and outlet gas compositions. The microGC was equipped with two columns, the Molecular Sieve 5A PLOT 0.25 mm, 10 m (Agilent, Santa Clara, United States) that used Argon as a carrier gas to measure $H_2$, $O_2$, $N_2$, $CH_4$, and CO, and the PoraPLOT Q PLOT, 0.25 mm, 10 m (Agilent, Santa Clara, United States) that used Helium as a carrier gas to measure $CO_2$, $N_2$ (combined with $O_2$), and $H_2S$. The microGC was calibrated before the run using six and four calibration levels for experiments 1 and 2, respectively (**Table S14, ESI†)**. Each level was sampled for four replicates, and the average was taken as the calibration point. The total method lasted 180 s with a sample time of 20 s. The injector and sample line temperatures were set at 110°C, and column temperatures and pressures at 60°C and 150 kPa, respectively.

#### 1.11 Biomass concentration analysis

The biomass correlation coefficient (K in g/L/$OD_{600}$), as defined in Valgepea*, et al.*[46], was found by sampling 60 mL from each bioreactor (6 Falcon™ tubes of 10 mL) at the end of each steady-state period. The samples were centrifuged, the supernatant removed, and two Falcon™ tubes combined with 0.5 mL of Millipore water in pre-weighed glass vials (548-0028; VWR International GmbH, Darmstadt, Germany), resulting in three technical replicates per bioreactor. The vials were dried at 200 mbar (absolute pressure) and 80°C for three days, and the weight of the biomass was recorded. The slope of the measured biomass weight to the corresponding $OD_{600}$ was taken and divided by the volume (0.02 L) to give the following K values (g/L/$OD_{600}$): First experiment: *M. thermautotrophicus* ΔH, 0.32; *M. thermautotrophicus* Z-245, 0.31; and *M. marburgensis* Marburg, 0.28. Second experiment: *M. thermautotrophicus* ΔH pMVS1111A:$P_{hmtB}$-$fdh_{Z\text{-}245}$ (molecular hydrogen and carbon dioxide), 0.31; *M. thermautotrophicus* ΔH pMVS1111A:$P_{hmtB}$-$fdh_{Z\text{-}245}$ (sodium formate), 0.89; and *M. thermautotrophicus* Z-245 (sodium formate), 0.95.

#### 1.12 Sodium formate concentration measurements

Sodium formate concentrations were analyzed *via* high-pressure liquid chromatography (HPLC) (SIL-40C, Shimadzu Europa, Duisburg, Germany) system that was equipped with an Aminex HPX-87H column (300 by 7.8 mm; Bio-Rad, CA, USA) and a refractive index detector (RID-20A). A 5 mM sulfuric acid solution was used as the eluent, with a flow rate of 0.6 mL min$^{-1}$ and a sample run time of 30-60 min. The oven temperature was set to 60°C, while the sample rack of the attached autosampler to 4°C. For HPLC sample preparation, all culture samples (0.5 mL ± 0.1 mL) were filtered using 0.22 µm filters (ROTILABO® PVDF, 13 mm, Carl Roth, Karlsruhe, Germany). Sodium formate calibration curves were prepared with concentrations ranging from 0.5-10 mM and 20-400 mM.

#### 1.13 Carbon and electron balance calculations

For the first experiment, initial carbon balances using molar flow rates ($\dot{n}$, mmol/h) were calculated according to conservation of mass (**Equation 3**) using the ideal gas law (**Equation 4**) with the following assumptions:

- The ideal gas constant of 8.3144 L·kPa/K/mol
- A temperature (T) of 25°C (298.15 K)
- A pressure (P) of 1 atm (101.325 kPa).

$$C_{in} - C_{out} = 0 \qquad \text{Equation 3}$$

$$CO_{2,in} - \left(CO_{2,gas\_out} + CO_{2,aq\_out} + CH_{4,gas\_out} + biomass_{aq\_out}\right) = 0$$

$$\dot{n} = \frac{P\dot{V}}{RT} \qquad \text{Equation 4}$$

The volumetric flow rate ($\dot{V}$, L/h) was found by multiplying the gas concentration (the fraction measured by the microGC) by the measured gas flow rate. This was done for the $CO_2$ of the inlet gas ($CO_{2,gas\_in}$), and the $CO_2$ ($CO_{2,gas\_out}$) and $CH_4$ ($CH_{4,gas\_out}$) of the outlet gas. In addition, Henry's law was used to account for the soluble $CO_2$ ($CO_{2,aq\_out}$) that was removed from the bioreactors in the effluent. To find the quantities of soluble bicarbonate and carbonate in the medium, the acid dissociation constants (and pKa) were found using the equations provided by Prieto and Millero[47] (**Data S3, ESI†**). The salinity of the medium (4.565 ppt), required for the equations in Prieto and Millero[47], was estimated by summing the amount of cations and anions of the salts that were added to the medium. The estimated concentrations of solubilized and removed carbon species were subtracted from the calculated gas uptake values. Lastly, to calculate the biomass concentration in the bioreactor ($biomass_{aq\_out}$), the biomass correlation coefficient (K in g/L/$OD_{600}$) was divided by the measured bioreactor $OD_{600}$ and volume. The resulting $g_{CDW}$ was divided by the molecular weight (23.502 g/mol) [44] and multiplied by the measured dilution rate of the bioreactors ($h^{-1}$). Given inconsistencies in carbon balances in our measurements, a gross measurement error analysis was performed following the macroscopic balance method described in Wang and Stephanopoulos[48] (**Data S6, ESI†**). In brief, first, the probable gross measurement error was identified *via* statistical hypothesis testing with a chi-square test. Second, maximum likelihood estimates were calculated (dropping the data points with gross measurement error) to find corrected values for the gas and biomass data.

For the second experiment, on molecular hydrogen and carbon dioxide, the carbon balances were performed as described above with an adjusted salinity value of 4.568 ppt rather than 4.565 ppt. However, given more considerable inconsistencies in carbon and electron balances with the raw data for bioreactor runs on sodium formate, a different method was applied in this case. No sodium formate was detected in the liquid media during steady-state. Thus, we assumed that the microbes consumed the entire sodium formate from the feed in the bioreactor ($Na\text{-}formate_{in}$). However, the quantity of 1 M HCl solution added to the bioreactors for pH control was not insignificant (unlike the amount of NaOH added for fermentations on molecular hydrogen and carbon dioxide). The feed rate of the pH control pumps was determined, and a dilution factor was calculated. Approximately 21.88 ± 2.93% of the liquid feed into the bioreactors was due to the acid feed, thus lowering the effective sodium formate feed concentration to 291.38 ± 10.55 mM (**Table S15, ESI†**). Further, the estimation of dissolved carbon species is very sensitive to salinity. The increased salt concentrations from the sodium formate (new value estimated at 17.80 ± 0.44 ppt) were insufficient for closing both carbon and electron balances. It is also noted that variation in pH was more frequent due to the production of NaOH with sodium formate consumption, and at times, there was a noticeable shift in the online pH probe measurement that required a manual offset after offline pH measurements. Furthermore, the dissolved carbon dioxide values are sensitive to changes in pH. Thus, to solve for the quantities of produced $CH_4$ and $CO_2$, **Equations 5** and **6**, which represent the carbon and electron (using degree of reduction) balances, were solved. The biomass coefficient in **Equation 6** was calculated assuming a molecular formula of $CH_{1.681}O_{0.481}N_{0.222}$, as described in Michael and Kargi[49]. Molecular hydrogen gas was detected in the outlet gas stream ($H_{2,gas\_out}$), and thus it was also included in **Equation 6.**

$$C_{in} - C_{out} = 0 \quad \text{Equation 5}$$

$$Na\text{-}formate_{in} - \left(CO_{2,gas\_out} + CO_{2,aq\_out} + CH_{4,gas\_out} + biomass_{aq\_out}\right) = 0$$

$$Na\text{-}formate_{in} - \left(CO_{2,gas+\quad\_out} + CH_{4,gas\_out} + biomass_{aq\_out}\right) = 0$$

$$E_{in} - E_{out} = 0 \quad \text{Equation 6}$$

$$2 * Na\text{-}formate_{in} - \left(8 * CH_{4,gas\_out} + 4.053 * biomass_{aq\_out} + 2 * H_{2,gas\_out}\right) = 0$$

#### 1.14 Normalized product distribution

Product distributions were calculated using $CH_4$ and biomass data (mmol/h) for bioreactors and time points that did not have suspected gross measurement errors (**Section 1.13, ESI†**). The quantity of $CH_4$ or biomass was divided by the sum of the $CH_4$ and biomass and then multiplied by 100, resulting in the normalized product distribution percentage. For fermentations on sodium formate, the carbon dioxide production was also accounted for in the products. An analysis of variance (ANOVA) test was conducted in Excel® using the `ANOVA:Single Factor` ($\alpha$ = 0.05) function of the Data Analysis Addin to analyze statistically significant differences between the product distribution ratios of the three different microbes (first experiment). Further, product distribution ratios were compared for: **1)** *M. thermautotrophicus* ΔH pMVS1111A:P$_{hmtB}$-*fdh*$_{Z\text{-}245}$ and *M. thermautotrophicus* Z-245 grown on sodium formate (second experiment); and **2)** product distribution ratios of *M. thermautotrophicus* ΔH, *M. thermautotrophicus* Z-245, and *M. marburgensis* Marburg grown on $H_2/CO_2$ (from the first experiment) with *M. thermautotrophicus* ΔH pMVS1111A:P$_{hmtB}$-*fdh*$_{Z\text{-}245}$ grown on $H_2/CO_2$ (from the second experiment).

##### 1.15 Interspecies comparison

For the first experiment, all gas and biomass data points without suspected gross measurement error were used for the comparative analysis (independent *t*-tests) between the three microbes, except for one bioreactor replicate for *M. thermautotrophicus* Z-245 that experienced a pump malfunction and wash-out of bioreactor broth during the fermentation.

For the second bioreactor experiment, one replicate of *M. thermautotrophicus* ΔH pMVS1111A:P$_{hmtB}$-*fdh*$_{Z\text{-}245}$ grown on sodium formate was discarded because of a mixer malfunction, and thus, the gas and biomass data points were not included in the *t*-tests. The rest of the gas and biomass data points were used for the comparative analysis between *M. thermautotrophicus* ΔH pMVS1111A:P$_{hmtB}$-*fdh*$_{Z\text{-}245}$ and *M. thermautotrophicus* Z-245 grown on sodium formate. The second set of *t*-tests was conducted between *M. thermautotrophicus* ΔH, *M. thermautotrophicus* Z-245, and *M. marburgensis* Marburg grown on $H_2/CO_2$ from the first experiment with *M. thermautotrophicus* ΔH pMVS1111A:P$_{hmtB}$-*fdh*$_{Z\text{-}245}$ grown on $H_2/CO_2$ from the second experiment.

The *t*-tests (two-tailed and confidence intervals of 95%) were carried out in Python 3.6.13 using the Researchpy package Version 0.3.2 and the `ttest` function.[50] Three-way comparisons were performed in Excel® using the `ANOVA:Single Factor` ($\alpha = 0.05$) function of the Data Analysis Addin. The maximum likelihood estimates of the measurements for time points without gross measurement error were calculated,[48] which resulted in closed carbon balances for those time points (assuming 90% confidence intervals for the test function). These adjusted values were then averaged across time points for each microbe and used to constrain the GEMs (**Data S3, ESI†**).

#### 1.16 RNA sample preparation

When the steady-state was reached, four technical replicates of 9 mL (*n*=4) of bioreactor sample were placed into 5 mL of prechilled RNAlater® (overnight at 4°C). The samples were stored overnight at 4°C and then frozen at -20°C until RNA isolation. The samples were thawed on ice and centrifuged at 4°C and 4100 × g for 10 min (Multifuge X3R TX-1000, Fischer Scientific, USA), the supernatant was discarded, and the samples were resuspended in 800 µL of RNase-free water. The 800 µL were mixed with 950 µL of saturated phenol (Sigma-Aldrich, Merck, Germany) and 115 µL of a lysis solution in Lysing Matrix B (MP Biomedicals Germany GmbH, Eschwege, Germany). The lysis solution contained sodium acetate (20 mM pH 5.2), sodium dodecyl sulfate (SDS; 0.5 % v/v), ethylenediaminetetraacetic acid (EDTA; 1 mM), and DNase/RNase-free distilled water (DI; Invitrogen, Thermo Fischer Scientific, Germany). The cells were homogenized for one cycle (5 x 40 s at 6 m/s and 20 s off) in a FastPrep-24™ 5G bead-beater (MP Biomedicals Germany GmbH, Eschwege, Germany) and then centrifuged at room temperature and 21130 × g for 10 min (5424 Eppendorf, Hamburg, Germany). The top layer was placed into 600 µL of phenol-chloroform-isoamyl alcohol (ROTI® Aqua-P/C/I, Carl Roth, Karlsruhe, Germany) and centrifuged as in the previous step. This step was repeated twice. The top layer was added to 1.2 mL of ethanol (undenatured absolute, SERVA Electrophoresis GmbH, Heidelberg, Germany) and left overnight at -80 °C to precipitate the RNA. The following day, the tubes were centrifuged at 4°C and 21130 × g for 10 min, and then the supernatant was removed. The RNA pellet was washed with 1 mL of 75% v/v ethanol solution and centrifuged as in the previous step. The supernatant was pipetted off, and the pellet was resuspended with 53 µL of DNase/RNase-free distilled water. The RNA was cleaned (which included DNA depletion) and concentrated using the RNA Clean & Concentrator kit (Zymo Research, Irvine, CA,

United States) according to the manufacturer's instructions. The cleaning and concentrating protocols were repeated four times. The quantity (6-16 µg) and quality (RNA integrity index > 9) of the purified RNA were measured with a 2100 Agilent Bioanalyzer (Agilent Technologies, Santa Clara, United States) using the Agilent RNA 6000 Nano Kit (Agilent Technologies, Santa Clara, United States) before freezing the samples at -80°C until sequencing.

#### 1.17 RNA sequencing

The library preparation was performed using 100 ng of RNA and the TruSeq™ Stranded Total RNA Kit with Ribo-Zero™ Plus (Illumina, San Diego, United States). Pair-ended sequencing was performed using the NovaSeq™ 6000 with the Flow Cell Type 2 x 100 bp (Illumina, San Diego, United States). Demultiplexing of the sequences was performed with Illumina bcl2fastq (2.20) software. The RNA sequencing was performed by CeGaT GmbH (Tübingen, Germany).

#### 1.18 Raw RNA sequencing data analysis

Sequencing reads were quality filtered and trimmed with BBMap version 38.93[51] and FastX Toolkit version 0.0.14[52] with specific parameters for paired-end reads. Reads from each bioreactor culture were mapped to the reference genomes of *M. thermautotrophicus* ΔH (`CP064324`), *M. thermautotrophicus* Z-245 (`CP064336` and `CP064337`), and *M. marburgensis* Marburg (`CP069376` and `CP069377`) using BWA-MEM version 0.7.17.[53] Mapped reads were assigned to genomic features using the FeatureCounts program from the Subread package version 2.0.1.[54] Additionally, an alignment-free quantification of gene features was performed using Salmon version 1.5.2.[55] Pairwise differential expression analysis was performed on the Salmon counts using DESeq2 version 1.32.0[56] with the homologous genes present in the pan-genome database (**Section 1.22, ESI†**) and ConsensusDE version 1.10.0.[57] The adjusted *P*-value cut-off threshold was set to 0.05. This process was orchestrated using Snakemake version 6.8.0[58] to achieve reproducibility (**Data S6, ESI†**). The output files from Salmon were used to analyze each microbe individually, and only genes with at least two non-zero transcript values were considered. The averages of each gene were then used.

#### 1.19 Protein sample preparation

The proteomics sampling and sample preparation methods from Valgepea*, et al.*[59] were used with the following modifications. Screw-cap 2-mL tubes with lysing matrix B (MP Biomedicals Germany GmbH, Eschwege, Germany) were used in the homogenizer (FastPrep-24TM 5G, MP Biomedicals Germany GmbH, Eschwege, Germany). For cell lysis, the one cycle of bead beating as described for transcriptomics sample preparation was repeated three times. The following protein precipitation method from the Proteome Center Tübingen (PCT) at the University of Tübingen was applied. The supernatant was transferred to a 15-mL FalconTM tube with 4 mL and 0.5 mL of ice-cold 100% acetone and 100% methanol, respectively, and precipitated overnight at -20°C. The following day, the tubes were centrifuged at 2200 × g and 4°C for 20 min (Multifuge X3R TX-1000, Thermo Fischer Scientific, Waltham, United States). The supernatant was removed, and the pellet was washed with 1 mL of ice-cold acetone (80/20% v/v in water) and centrifuged as in the previous step. The supernatant was removed, and the protein pellet was left to air dry on ice for 15 min before freezing at -20°C.

#### 1.20 Protein measurement

Pellets were resuspended in a denaturation buffer (6 M urea, 2 M thiourea in 10 mM Tris pH 8.0), and 20 µg of protein were subjected to tryptic in-solution digestion (0.2 µg trypsin). The samples were run for an LC-MS/MS analysis on an Easy-nLCTM 1200 (Thermo Fisher Scientific, Dreieich, Germany) coupled to an Orbitrap Exploris™ 480 mass spectrometer (Thermo Fisher Scientific, Dreieich, Germany). The LC-MS analysis was performed as described previously in Fagbadebo*, et al.*[60], with the exception of the duration and gradient of the peptide elution. Solvent A was 0.1% formic acid, and solvent B was 80% acetonitrile in 0.1% formic acid [60]. The LC-MS was operated at 40 °C.[60] The peptides were eluted from 0-113 min with a linear gradient from 10% to 33% of solvent B at a flow rate of 200 nL/min.

Then, from 113-116 minutes, the gradient increased from 33% to 50% of solvent B at 200 nL/min. To wash the remaining peptides from the column, from minutes 116-119, the gradient increased from 50% to 90% of solvent B with a gradually increasing flow rate from 200 nL/min to 500 nL/min. Lastly, the column was then washed from 119-127 min with 90% solvent B at 500 nL/min.

##### 1.21 Protein analysis with proteome discoverer

Raw data-dependent acquisition (DDA) data from LC-MS/MS analysis was performed using Thermo Fisher Proteome Discoverer software (version 2.5.0.400, Thermo Fisher Scientific, Scoresby, Australia). Files were processed using two different methods: **1)** for intensity-based absolute quantification (iBAQ) analysis, each strain was processed individually through the precursor label-free quantification workflow with their corresponding FASTA files and then converted to iBAQ as described in Schwanhäusser, *et al.*[61]; and **2)** for differential expression analysis, all files were processed together using the custom pan-genome FASTA file (**Section 1.22, ESI†**). Both methods used the same workflows and only differed in their choice of FASTA file. Briefly, spectra were searched against the respective databases with carbamidomethylation set as a fixed modification and up to three methionine oxidations. Acetylation, methionine loss, and acetylation after methionine loss at the N-terminus were also permitted. The mass tolerance of precursors and fragments were 10 ppm and 0.02 Da, respectively. The minimum and maximum peptide lengths were six and 144 amino acids, respectively. Two missed cleavages were allowed per peptide. Proteins were filtered to a 1% FDR cut-off threshold and for an adjusted *P*-Value of 0.05. For quantification, only peptides unique to a protein group were used. All other parameters were kept at their default settings.

##### 1.22 Pan-genome differential expression database creation

With the *M. thermautotrophicus* $\Delta$H genes as a reference, an intersection of homologous genes/proteins based on the best BLASTp+ hits from *M. thermautotrophicus* Z-245 and *M. marburgensis* Marburg was used (**Equation 7**).

*M. thermautotrophicus* ΔH and *M. thermautotrophicus* Z-245 had 81 additional homologous genes/proteins, which were not considered when *M. marburgensis* Marburg was being analyzed, while *M. thermautotrophicus* ΔH and *M. marburgensis* Marburg had 11. This pan-genome of homologous gene groups can be found in **Data S1, ESI†**.

$$(ZZ \cup MM) \cap DH \qquad \text{Equation 7.}$$

A protein FASTA file that represented the homologous clusters was required to perform relative proteomics across different species. Each homologous cluster FASTA entry needs to account for genetic heterogeneity that leads to variation in protein sequences across microbes. Therefore, peptide sequences from the other microbes had to be added to the gene-group sets.[62] An *in-silico* tryptic digestion of the homologous protein FASTA sequences for *M. thermautotrophicus* ΔH, *M. thermautotrophicus* Z-245, and *M. marburgensis* Marburg was performed using Rapid Peptides Generator version 1.2.4 to retrieve all peptides for each microbe.[63] A new enzyme was defined to cleave after lysine (K) or arginine (R) except if proline (P) follows, with the cleaving rule as `(K or R,)` with the exception `(K or R,)(P)`. Then, using the *M. thermautotrophicus* ΔH protein FASTA sequence as a reference, unique peptides (with a length greater than six amino acids) from homologous proteins of *M. thermautotrophicus* Z-245 and *M. marburgensis* Marburg that were not already in the *M. thermautotrophicus* ΔH protein were inserted *prior* to the C-terminal peptide (**Data S6, ESI†**). If *M. thermautotrophicus* Z-245 and *M. marburgensis* Marburg shared a C-terminus different from the one of *M. thermautotrophicus* ΔH, then that C-terminus was additionally amended as the C-terminus for the new combined protein, otherwise the C-terminus from *M. thermautotrophicus* ΔH was kept. This new protein FASTA file was then used for the proteomics analysis (**Section 1.21, ESI†).**

#### 1.23 Eha/Ehb ratio determination

The ratio of the energy-converting hydrogenases Eha and Ehb subunits was found by summing the abundances (either transcripts or relative protein abundances) of the subunits detected for each energy-converting hydrogenase and dividing those sums by the number of subunits found. This was done to normalize against subunits that were not found (the stoichiometry between all subunits was considered as equimolar based on current knowledge from the literature).[64] The normalized abundance of Eha was then divided by that of Ehb. This ratio was then averaged across the replicates (bioreactor samples) of each microbe. An ANOVA test was conducted in Excel® using the `ANOVA:Single Factor` ($\alpha = 0.05$) function of the Data Analysis Addin to test if there was a statistically significant difference between the Eha/Ehb ratios for the three different microbes (**Data S1, ESI†**).

##### 1.24 Methanogenesis relative abundances

Gap-filled relative abundances were found for the different proteins (or complexes) involved in the Wolfe cycle of methanogenesis. The iBAQ values of each subunit were summed and divided by the number of detected subunits (the maximum number of subunits missing per microbe per protein complex was one). This sum was then multiplied by the theoretical number of subunits that should be found (effectively assigning the non-detected subunit the average iBAQ value for the complex) to produce the entire protein complex normalized iBAQ value. This value was then divided by the sum of all iBAQ values in the proteome and multiplied by 100, resulting in relative (normalized) protein abundance percentages (**Data S1, ESI†**). The stoichiometry between all subunits was considered as equimolar based on current knowledge from the literature.[64]

##### 1.25 Integrating fermentation data in the GEMs

Experimental fermentation data (Conditions 22-24) were used to constrain the models for the FBAs (with biomass maximization as the objective function) to validate the three strain-specific GEMs (**Data S4 and Data S6, ESI†**). The data did not include time points with suspected gross measurement error and was adjusted with the maximum likelihood estimates.

Reduced experimentally-constrained GEMs were created by applying the GIMME[65] algorithm with the transcriptomics and proteomics data (thresholds set at the lower quartile). This step was accomplished in the COBRA Toolbox v.3.1[66] using MATLAB (R2018b) and the Gurobi Optimizer v.9.0.1[67]. Also in the COBRA Toolbox, FBAs (with and without loops allowed) were run on the resulting reduced GEMs, using either the maximization of biomass exchange (`EX_biomass_e`) or the maximization of ATP dissipation (`ATPM`) as the objective function (**Data S4 and Data S6, ESI†**).

### 2 Results

#### 2.1 Supplementary Text

##### Text S1. Methylation patterns of the microbes

The methylation patterns of the three microbes were elucidated by single-molecule real-time long-read sequencing (**Sections 1.2-1.3, ESI†***)*. The microbes exhibited N6-methyladenine (m6A) and N4-methylcytosine (m4C) methylation in three distinct methylation patterns **(Table S2, ESI†**). The more closely related *M. thermautotrophicus* ΔH and *M. thermautotrophicus* Z-245 shared one sequence motif (with an m6A modification) that exhibited features of a Type-I restriction/modification system(**Table S1, ESI†**).[68] The two m6A methylation patterns from *M. marburgensis* Marburg were more distinct but also characteristic of Type-I restriction/modification systems (**Table S1, ESI†**). The genomes of all three microbes encode putative components of Type-I restriction/modification systems (**Table S1, ESI†**), which could be responsible for the methylation of the DNA.[69,70] *M. marburgensis* Marburg had an additional m4C modification, which resembles a Type-II modification, with putative methyl-transferase genes encoded in the genome (**Table S1, ESI†**). *M. thermautotrophicus* Z-245 also had an additional motif with an m4C modification for which the responsible Type-II restriction/modification methyl-transferase is located on the plasmid (pFZ1),[71] assigning a known function to the plasmid. While *M. marburgensis* Marburg also harbors a plasmid (pME2001), its function is still unknown but does not encode obvious restriction/modification systems.

**Text S2. Multi-level omics under steady-state growth conditions indicate elevated oxidative stress response**

We did not find apparent reasons for the different growth behaviors when analyzing methanogenesis gene products. In the multi-level omics analysis, we found several genes and gene products that are potentially involved in oxidative stress response and were highest on average in *M. marburgensis* Marburg compared to the other two microbes. These included, in the transcriptomics (though not on the proteomics level), the $F_{390}$ synthase (gene group 1317) that protects cofactor $F_{420}$ (and halts methanogenesis) by converting it into the oxygen-protected form, cofactor $F_{390}$,[72] and in the proteomics (though not on the transcriptomics level), three enzymes (gene groups 128, 186, and 1155) that are annotated as $F_{420}H_2$-dependent oxidases that convert dioxygen into water. While we measured dioxygen in the range of 0.069 ± 0.085% v/v in the exhaust gas of all bioreactors, indeed, we found significantly higher dioxygen levels of 0.315 ± 0.098 (*P*-value 7.38E-77) in the exhaust gas of one of the bioreactors with *M. marburgensis* Marburg (**Figure S5, ESI†**). Thus, our results confirmed our observation of increased dioxygen levels in an increased level of oxidative stress in the averaged transcriptome and proteome in *M. marburgensis* Marburg.

**Text S3. Prediction of the conversion of methenyl-H4MPT to methylene-H4MPT with integration of multi-omics data**

In all three microbes, we found a high abundance of the enzymes $F_{420}$-dependent methylene-$H_4$MPT dehydrogenase (Mtd, reaction MTD), $H_2$-forming methylene-$H_4$MPT dehydrogenase (Hmd, reaction HMD), and $F_{420}$-reducing Ni-Fe hydrogenase (Frh, reaction FRH), which are required to catalyze the conversion of methenyl-$H_4$MPT to methylene-$H_4$MPT (**Figure 3**). Frh generates reduced cofactor $F_{420}$ and, thus, provides reducing equivalents for Mtd under standard conditions. During nickel limitation, the conversion of methenyl-$H_4$MPT to methylene-$H_4$MPT is catalyzed by Mtd together with Hmd instead, also known as the Hmd-Mtd cycle.[73] Under these conditions, the complex of Mtd and Hmd can also replace Frh to provide reduced cofactor $F_{420}$ by catalyzing the reaction in reverse.[73,74] While the Frh was found to be experimentally reversible *in vitro*,[74] we assumed irreversibility for our modeling in the direction of reduced cofactor $F_{420}$ production because molecular hydrogen was present in excess in our experiments. The presence of all three proteins (including Hmd) in the transcriptomics and proteomics data indicates that there might be a slight nickel-limitation, potentially occurring after specific biomass concentrations were reached (**Data S1, Data S3, and Data S4, ESI†**), which resulted in the partial bypassing of FRH in our modeling results. Alternatively, the presence of Hmd could be an example of the evolutionary tradeoff between rapid response time to changing environmental conditions (*e.g.,* limited nickel) and lowering *de-novo* protein synthesis costs.[75] Although all three enzymes were present, constraining the GEMs with the omics data did not result in a solution that uses all three reactions simultaneously (unless FRH was set to work reversibly, which generated a loop with the other two reactions). Surprisingly, the omics-constrained flux balance analyses predicted the use of HMD with twice the flux and MTD working in reverse for all simulations, with no flux carried by FRH. Exceptions were found for the transcriptomics-reduced *M. thermautotrophicus* ΔH GEM (Simulations 29-32), the transcriptomics-reduced *M. thermautotrophicus* Z-245 GEM (Simulations 41-44), and the loopless *M. marburgensis* Marburg GEM (Simulation 51) for which HMD and FRH were both active in the forward direction (**Data S4, ESI†**). It should be noted that using a different algorithm to constrain the GEMs with the transcriptomics and proteomics data may result in alternative reactions carrying flux and different flux values.[76,77] However, our modeling results need to be further experimentally validated to resolve the fluxes that are carried in the MTD, HMD, and FRH reactions.

**Text S4. Predicted pyruvate-formate lyase-activating enzymes are present and highly upregulated in *M. thermautotrophicus* Z-245 and *M. marburgensis* Marburg despite the missing pyruvate formate-lyase main subunit.**

We only identified pyruvate formate-lyase in *M. thermautotrophicus* ΔH. An additional activating radical SAM enzyme, pyruvate formate-lyase 2 activating enzyme (gene group 279) that is present in *M. thermautotrophicus* ΔH (MTH345/ISG35_1595),[74,78] also had low confidence BLASTp+ hits for *M. thermautotrophicus* Z-245 and *M. marburgensis* Marburg. In addition, *M. thermautotrophicus* ΔH had low confidence annotations for two additional Pfl-activating enzymes (gene groups 917 and 1372), which also contain an annotated radical SAM core domain (Uniprot, MTH1069 and MTH1586). In turn, these two putative Pfl-activating enzymes had low confidence BLASTp+ hits for *M. thermautotrophicus* Z-245 and *M. marburgensis* Marburg.

The differential transcriptomics analysis showed upregulation of the gene that encodes the activating protein in gene group 279 in *M. thermautotrophicus* ΔH compared to the other two microbes (**Data S1, ESI†**). However, the opposite was found for the other putative activating enzymes (gene groups 917 and 1372), which were highly upregulated in *M. thermautotrophicus* Z-245 ($\log_2$FC of 5) and *M. marburgensis* Marburg ($\log_2$FC of 6). The upregulation of these gene groups may indicate their importance to other glycyl free radical (*e.g.,* SAM)-dependent enzymes whose function is still unknown (**Data S1, ESI†**).[78]

#### 2.2 Supplementary Figures

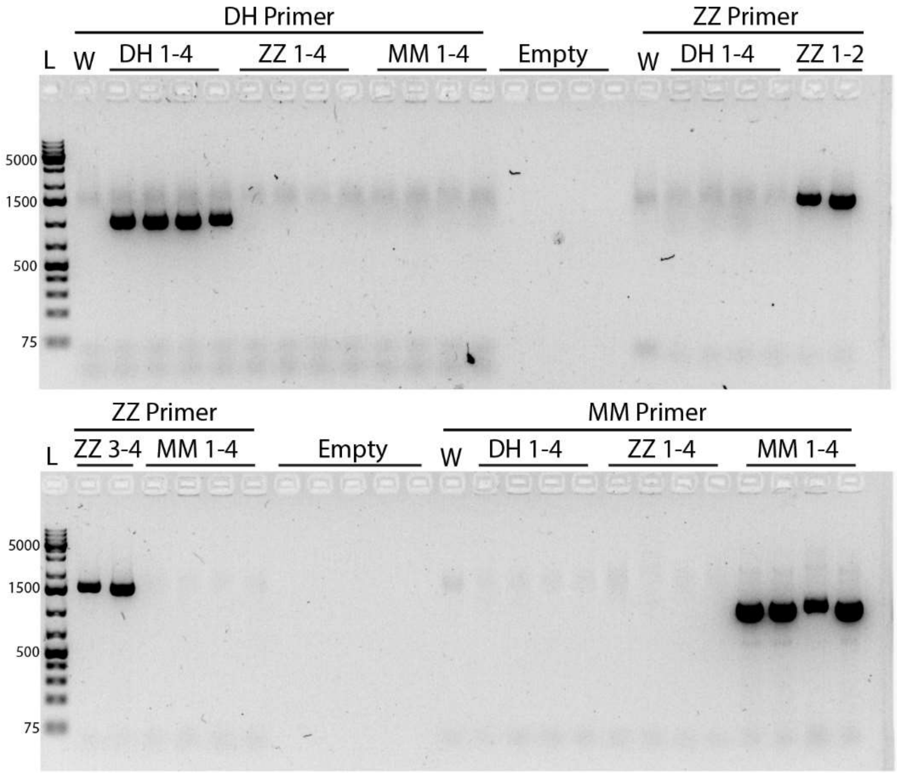


**Figure S1.** Check for cross-contamination in the bioreactors during Period 2 of Fermentation 1. L, ladder (1kbp); W, Millipore water; DH, *M. thermautotrophicus* ΔH; ZZ, *M. thermautotrophicus* Z-245; MM, *M. marburgensis* Marburg. Primers are listed in **Table S13, ESI†**.

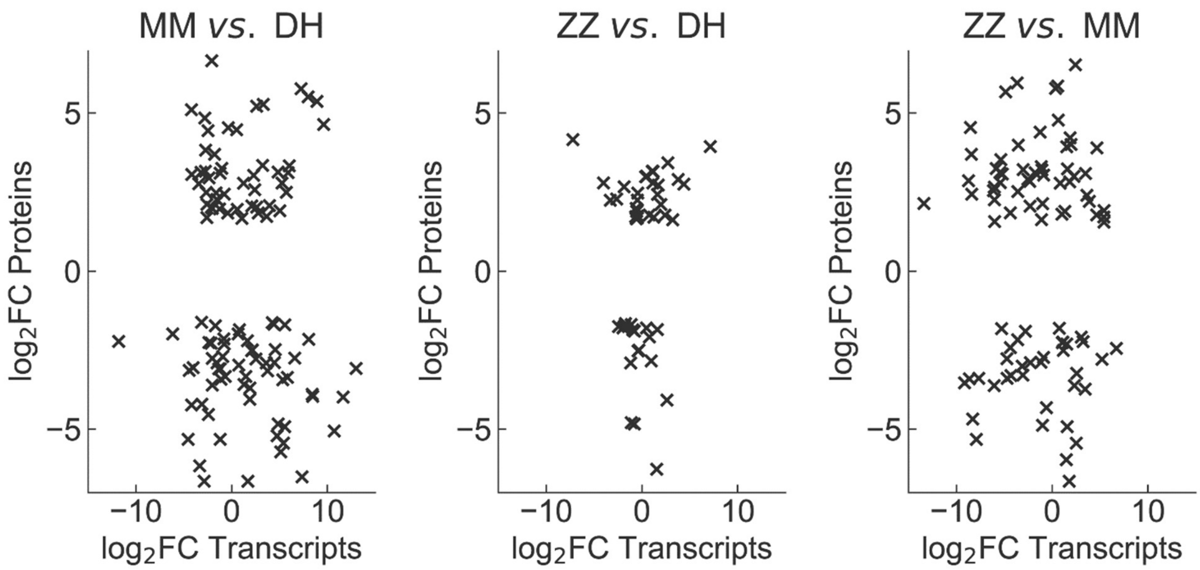


**Figure S2**. Significant (adjusted *P*-value ≤ 0.05) differentially abundant proteins versus differentially expressed genes (transcripts) for the three pairwise comparisons. DH, *M. thermautotrophicus* ΔH; ZZ, *M. thermautotrophicus* Z-245; MM, *M. marburgensis* Marburg

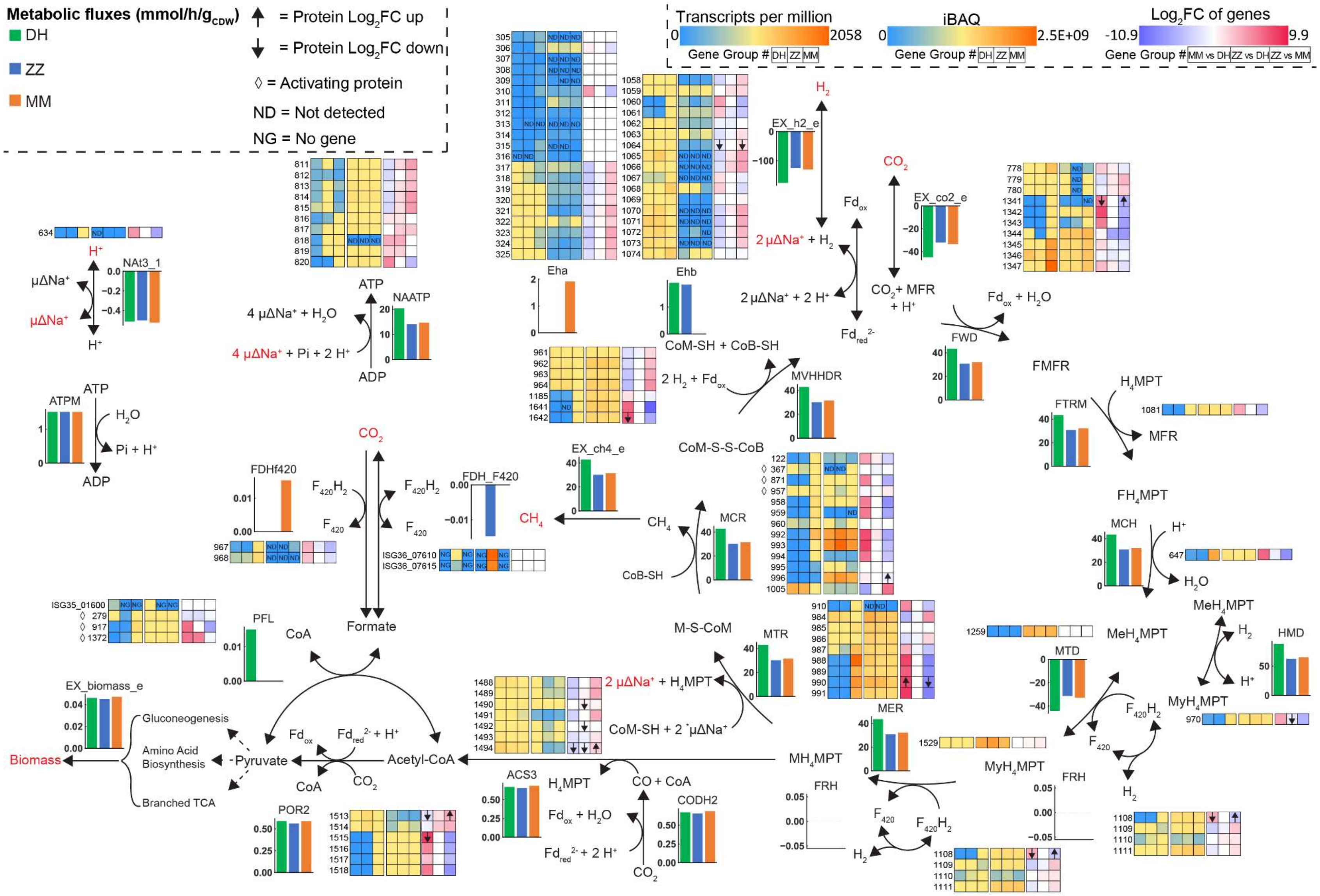

Metabolic fluxes (mmol/h/gCDW)
DH
ZZ
MM
= Protein Log2FC up
= Protein Log2FC down
◊ = Activating protein
ND = Not detected
NG = No gene
Transcripts per million
0
2058
Gene Group # DH ZZ MM
iBAQ
0
2.5E+09
Log2FC of genes
-10.9
9.9
Gene Group # MM vs DH ZZ vs DH ZZ vs MM
NAt3_1
ATPM
NAATP
ATP
ADP
H2O
Pi + H+
4 μΔNa+ + H2O
4 μΔNa+ + Pi + 2 H+
Eha
Ehb
H2
EX_h2_e
2 μΔNa+ + H2
2 μΔNa+ + 2 H+
Fdox
Fdred2-
CO2
EX_co2_e
CO2 + MFR + H+
Fdox + H2O
FWD
FMFR
H4MPT
FTRM
MFR
FH4MPT
MCH
H+
H2O
MeH4MPT
HMD
MyH4MPT
MTD
F420H2
F420
MVHHDR
CoM-SH + CoB-SH
2 H2 + Fdox
CoM-S-S-CoB
EX_ch4_e
CH4
MCR
CoB-SH
FDHf420
FDH_F420
Formate
CoA
PFL
M-S-CoM
MTR
2 μΔNa+ + H4MPT
CoM-SH + 2 μΔNa+
MER
MH4MPT
FRH
EX_biomass_e
Biomass
Gluconeogenesis
Amino Acid Biosynthesis
Branched TCA
Pyruvate
Acetyl-CoA
POR2
ACS3
CO + CoA
CODH2
Fdox + H2O
Fdred2- + 2 H+

**Figure S3**. Branched Wolfe Cycle adapted from Thauer[79] with other reactions involved in the energy metabolism. Fluxes are from the proteomics reduced model (using iBAQ values), constrained with experimental data that was adjusted for gross measurement error. Transcripts per million (TPM) for genes and iBAQ values for proteins. $Log_2$ fold change ($log_2FC$) for differentially expressed genes and proteins. Gene group is used as ID for the omics. For the PFL reaction, only *M. thermautotrophicus* ΔH has the gene; thus, the gene ID is used (`ISG35_01600`). For the FDH_F420, only *M. thermautotrophicus* Z-245 has the formate dehydrogenase cassette; thus, the gene IDs are used (`ISG36_07610` and `ISG36_07615`). Red text refers to metabolites that are exchanged across the membrane. **Microbes**: DH, *M. thermautotrophicus* ΔH; MM, *M. marburgensis* Marburg; ZZ, *M. thermautotrophicus* Z-245. **Compounds**: $CH_4$, methane; CO, carbon monoxide; CoA, Coenzyme A; CoB, coenzyme B; CoM, coenzyme M; CoM-S-S-CoB, CoM-CoB heterodisulfide; $CO_2$, carbon dioxide; F, formyl; $Fd_{ox/rd}$, ferredoxin oxidized/reduced; $H_2$, hydrogen; $H^+$, proton; $H_4MPT$, tetrahydromethanopterin; M, methyl; Me, methenyl; MFR, methanofuran; My, methylene; $Na^+$, sodium ion. **Reactions/Enzymes:** `ATPM`, ATP maintenance (pseudo reaction); `CODHr2`, CO dehydrogenase/acetyl-CoA synthase; `Eha/Ehb`, energy converting hydrogenases; `EX_biomass_e`, biomass exchange (pseudo reaction); `EX_ch4_e`, $CH_4$ exchange (pseudo reaction); `EX_co2_e`, $CO_2$ exchange (pseudo reaction); `EX_h2_e`, $H_2$ exchange (pseudo reaction); `FDHf420`, $F_{420}$-dependent formate dehydrogenase; `FDH_F420`, $F_{420}$-dependent formate dehydrogenase cassette; `FRH`, $F_{420}$-reducing hydrogenase; `FTRM`, FMFR/$H_4$MPT formyltransferase; `FWD`, FMFR dehydrogenase (tungsten- and molybdenum-dependent isozymes); `HMD`, Me$H_4$MPT hydrogenase; `MCH`, Me$H_4$MPT cyclohydrolase; `MCR`, MCoM reductase (I and II); `MER`, My$H_4$MPT reductase; `MTD`, My$H_4$MPT dehydrogenase; `MTR`, M$H_4$MPT/CoM methyltransferase; `MVHHDR`, $F_{420}$-non-reducing hydrogenase with the heterodisulfide reductase; `NAATP`, ATP synthase; `Nat3_1`, $Na^+/H^+$ antiporter; `PFL`, pyruvate formate-lyase; `POR2`, pyruvate synthase. **Other:** ◊, activating protein; ND, not detected; NG, no gene.

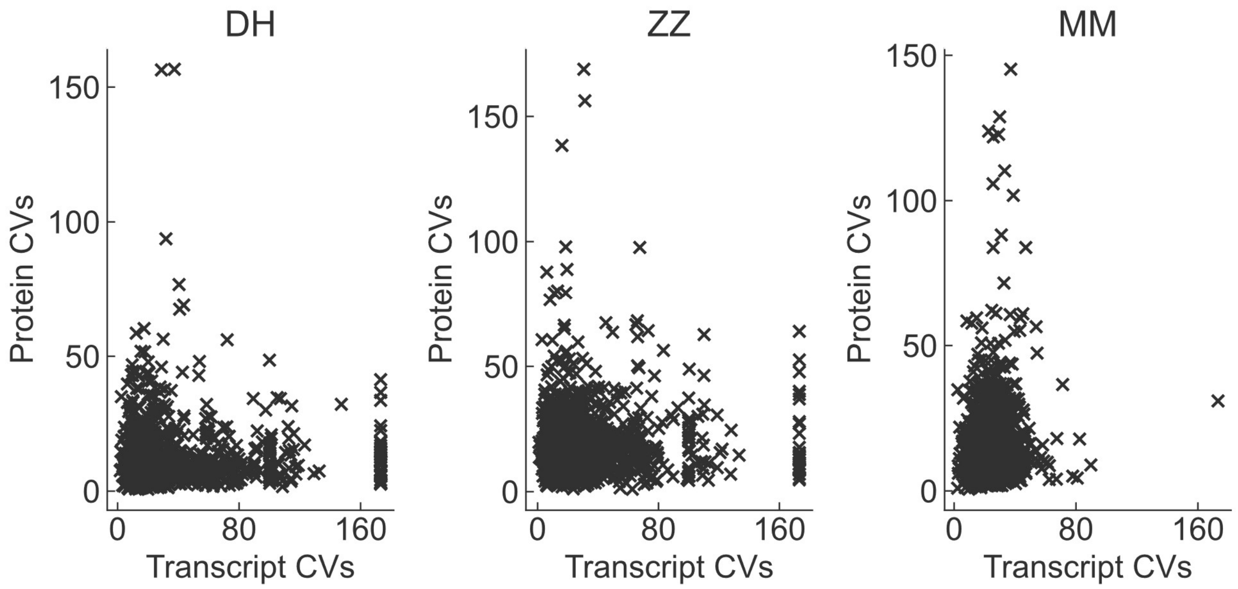


**Figure S4**. Coefficient of variance (CV) values of the proteins (proteome) versus the transcripts (transcriptome) of *M. thermautotrophicus* ΔH, *M. thermautotrophicus* Z-245, and *M. marburgensis* Marburg. DH, *M. thermautotrophicus* ΔH; ZZ, *M. thermautotrophicus* Z-245; MM, *M. marburgensis* Marburg

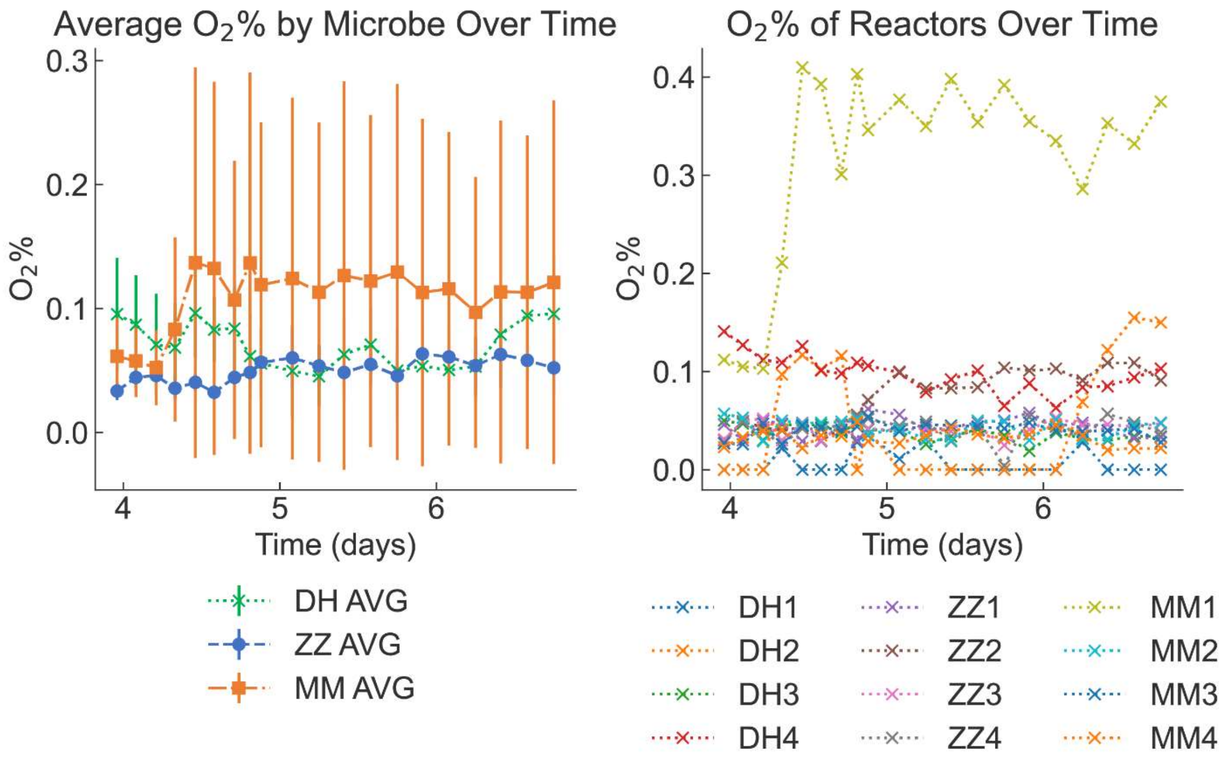


**Figure S5**. Dioxygen ($O_2$) levels (% v/v) in the bioreactor exhaust lines during Period 2 of Fermentation 1, averaged by microbe and by each bioreactor

replicate. DH, *M. thermautotrophicus* ΔH; ZZ, *M. thermautotrophicus* Z-245; MM, *M. marburgensis* Marburg.

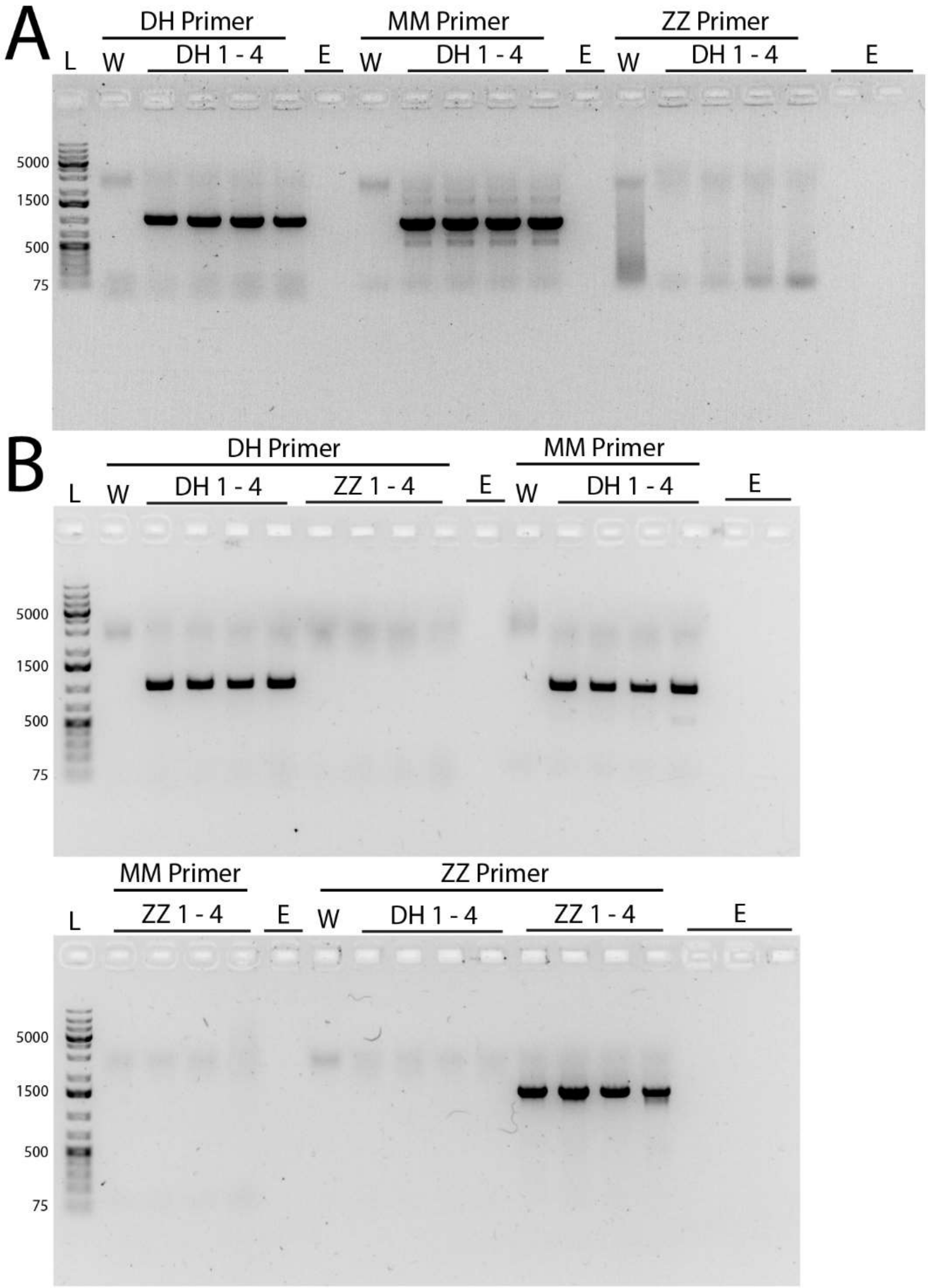


**Figure S6.** Check for cross-contamination in the bioreactors during steady-state of the second experiment. **A)** *M. thermautotrophicus* ΔH pMVS1111AP$_{hmtB}$-*fdh*$_{Z\text{-}245}$ cultivated on molecular hydrogen and carbon dioxide; **B)** *M. thermautotrophicus* ΔH pMVS1111AP$_{hmtB}$-*fdh*$_{Z\text{-}245}$ and *M. thermautotrophicus* Z-245 cultivated on sodium formate. E, empty lane; L, ladder (1kbp); W, Millipore water; DH, *M. thermautotrophicus* ΔH pMVS1111AP*$_{hmtB}$*-*fdh*$_{Z-245}$; ZZ, *M. thermautotrophicus* Z-245; MM, *M. marburgensis* Marburg. Primers are listed in **Table S13, ESI†**.

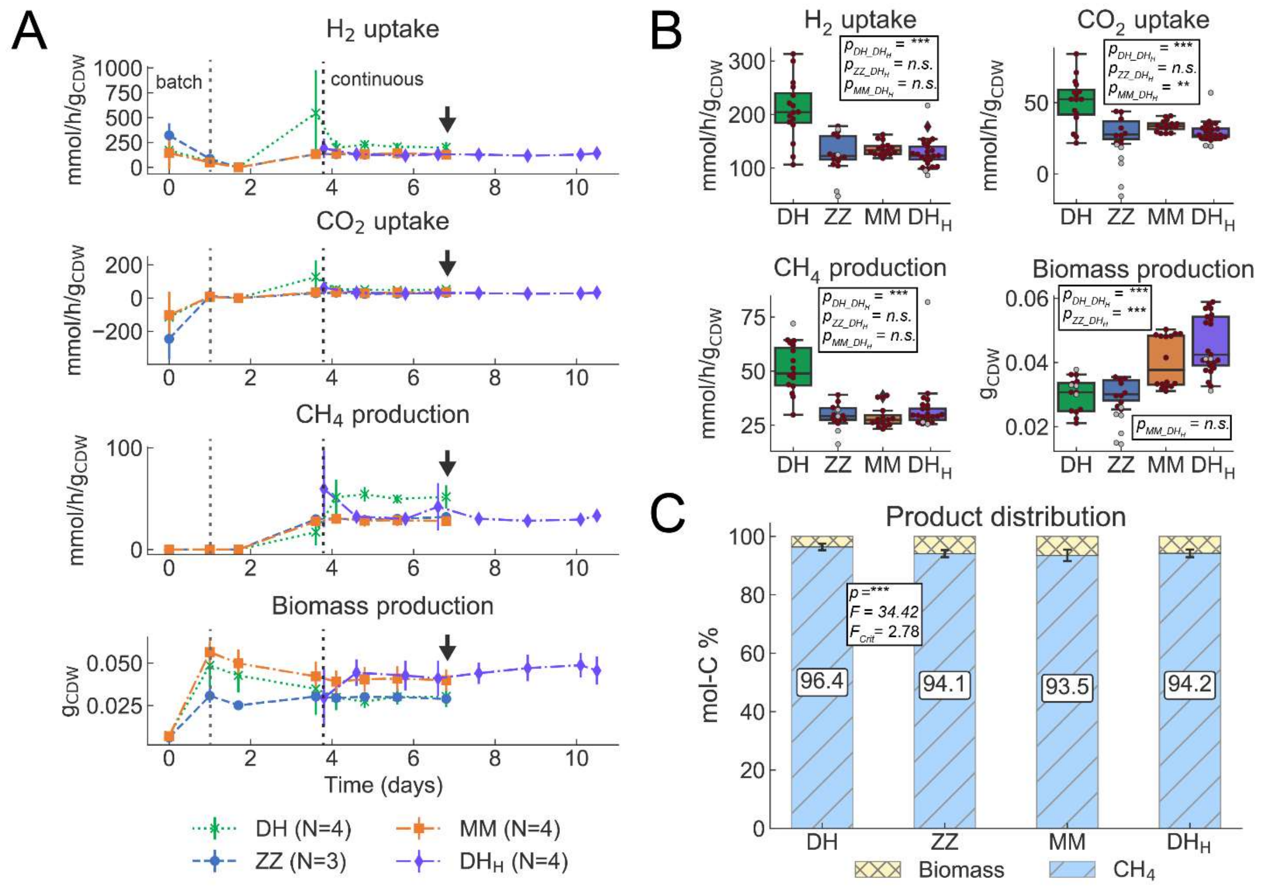


**Figure S7**. Fermentation data from chemostat bioreactors with *M. thermautotrophicus* ΔH with plasmid (pMVS1111AP$_{hmtB}$-fdh$_{Z-245}$) grown on $H_2$ and $CO_2$ ($DH_H$) in the second experiment, with data from the first experiment of *M. thermautotrophicus* ΔH (DH), *M. thermautotrophicus* Z-245 (ZZ), and *M. marburgensis* Marburg (MM) also grown on $H_2$ and $CO_2$. **A)** Gas consumption ($H_2$ and $CO_2$ uptake), and $CH_4$ and biomass production data from quadruplicate (DH, MM, and $DH_H$) and triplicate (ZZ) bioreactors for the fermentation period of 7 (DH, ZZ, MM) or 11 ($DH_H$) days. Data for further analyses (transcriptomics, proteomics) were taken on day seven as indicated by black arrows (DH, ZZ, MM). **B)** Average gas consumption ($H_2$ and $CO_2$ uptake) and $CH_4$ and biomass production data during a steady-state period (days 4 to 7 (DH, ZZ, MM) and days 5 to 11 ($DH_H$)). For statistical analysis in pairwise comparisons with *t*-tests, data points without suspected gross measurement error (red circles) were included, and data points with suspected gross measurement error (gray circles) were excluded (**Section 1.13, ESI†**). **C)** Average normalized product distribution, including statistical analysis by ANOVA (**Section 1.14, ESI†)**. DH, *M. thermautotrophicus* ΔH (without plasmid); $DH_H$, *M. thermautotrophicus* ΔH harboring $pMVS1111AP_{hmtB}$-$fdh_{Z\text{-}245}$; MM, *M. marburgensis* Marburg; ZZ, *M. thermautotrophicus* Z-245; ***, $p < 0.0001$; **, $p < 0.001$; n.s., not significant ($p > 0.05$); $F$, $F$ value; $F_{crit}$, $F$ critical value.

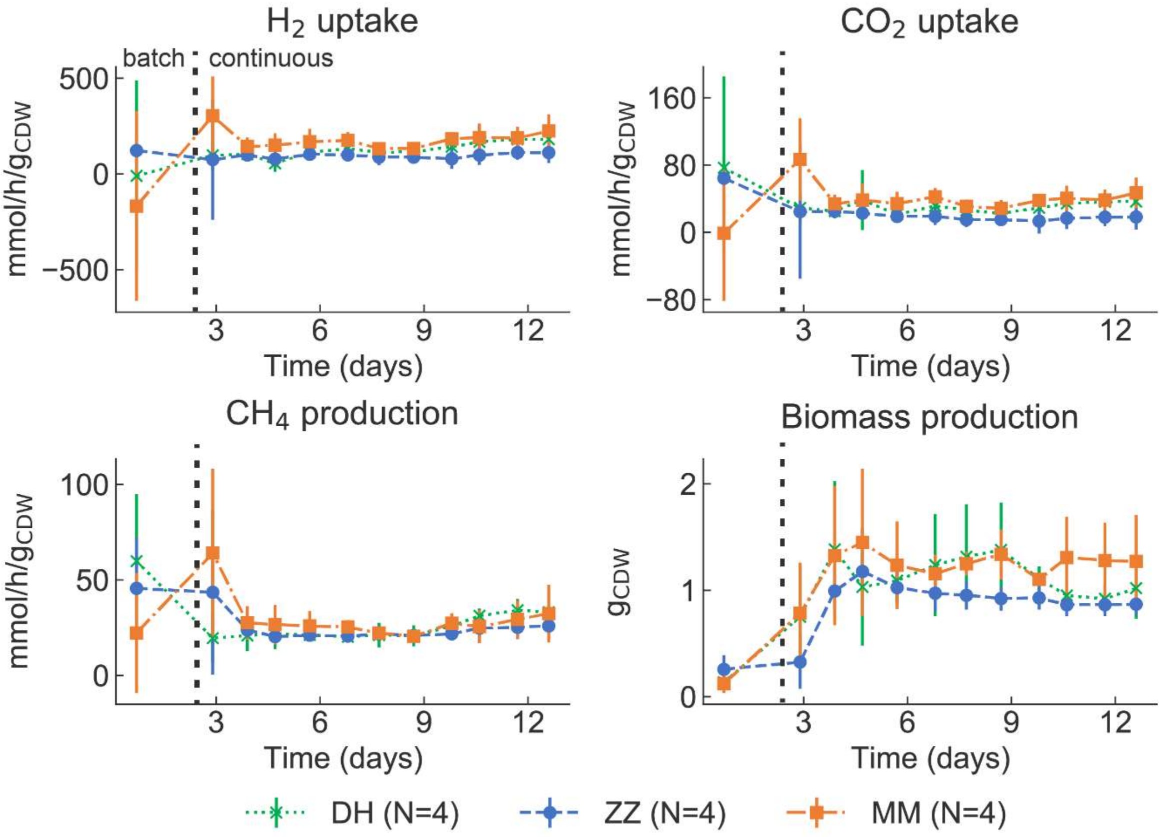


**Figure S8**. Gas consumption ($H_2$ and $CO_2$ uptake), and $CH_4$ and biomass production data from quadruplicate bioreactors for the fermentation period of 13 days (first experiment, period 1). DH, *M. thermautotrophicus* ΔH; ZZ, *M. thermautotrophicus* Z-245; MM, *M. marburgensis* Marburg.

#### 2.3 Supplementary Tables Legends

*Tables are found in tables.xlsx, ESI†*

**Table S1**. Details on access to genome assembly information in NCBI.

**Table S2**. Methylation patterns of the three microbes elucidated through PacBio sequencing

**Table S3**. COG categorization of the genomes of the following microbes: DH, *M. thermautotrophicus* ΔH; ZZ, *M. thermautotrophicus* Z-245; and MM, *M. marburgensis* Marburg

**Table S4**. Differences between new and old genome sequences in *M. thermautotrophicus* ΔH and *M. marburgensis* Marburg

**Table S5**. Summary of fermentation production rates

**Table S6**. Differential protein expression analysis of proteins that mapped to reactions in the model

**Table S7**. Genes in top 100 most abundant transcripts that are responsible for reactions in the model

**Table S8**. Proteins in top 100 most abundant gene products (based on averaged iBAQ values) that are responsible for reactions in the model

**Table S9**. Percent of average relative abundance of proteins (of proteome) involved in methanogenesis (based on iBAQ values) normalized to the number of subunits found to how many would be present (each * indicates one subunit was missing)

**Table S10**. Ratios of Eha to Ehb based on the averages within each microbe (normalized to the number of subunits detected)

**Table S11**. Number of genes related to biomass synthesis either up and down regulated in MM to the other microbes (from the transcriptomics study): DH, *M. thermautotrophicus* ΔH; ZZ, *M. thermautotrophicus* Z-245; and MM, *M. marburgensis* Marburg

**Table S12**. Tools used for the genome assembly and methylation pattern detection: DH, *M. thermautotrophicus* ΔH; ZZ, *M. thermautotrophicus* Z-245; and MM, *M. marburgensis* Marburg

**Table S13**. Microbe specific primers. The primers for *M. thermautotrophicus* Z-245 and *M. marburgensis* Marburg bind on the respective plasmids

**Table S14**. Calibration gas levels for Agilent 490 MicroGC, red values are not used in the calibration curve

**Table S15**. Dilution of continuous media (including adjusted sodium formate concentrations) based on base and acid feeds

#### 2.4 Data Files Descriptions

*Data files S1 and S3-S5 are found in Data_SX.zip, ESI†. Data files S2 and S6 are online resources, and their descriptions are also listed below.* DH, *M. thermautotrophicus* ΔH; ZZ, *M. thermautotrophicus* Z-245; and MM, *M. marburgensis* Marburg

**Data S1**: Excel® workbook with omics data

- Sheets 1-3: Transcriptomics individual DH, ZZ, MM
- Sheets 4-6: Transcriptomics differential expression MM_DH, ZZ_DH, ZZ_MM
- Sheets 7-9: Proteomics individual DH, ZZ, MM
- Sheets 10-12: Proteomics differential expression MM_DH, ZZ_DH, ZZ_MM
- Sheet 13: Relative abundances of Wolfe Cycle transcripts/proteins (iBAQ)
- Sheet 14: Relative abundances of Eha/Ehb transcripts/proteins (iBAQ)
- Sheet 15: Pan-genome

**Data S2**: BioModels links to the modeling files:
https://www.ebi.ac.uk/biomodels/MODEL2211290001 (*i*MTD22IC, DH)
https://www.ebi.ac.uk/biomodels/MODEL2211290002 (*i*MTZ22IC, ZZ)
https://www.ebi.ac.uk/biomodels/MODEL2211290001 (*i*MMM22IC, MM)

Each link contains a main model `.omex` file (which includes files listed below), an unbound FROG analysis `.omex` file generated using the unbound `.xml` model file, and a bound FROG analysis `.omex` file generated using the bound `.xml` model file.

- File 1: Pan-model Excel® model workbook (`.xlsx`)
- File 2: Pan-model biomass equation calculation Excel® (`.xlsx`)
- File 3: unbound `.xml` model files (CobraPy/Python work)
- File 4: unbound `.json` model files (Escher work)
- File 5: unbound `.mat` model files (COBRA Toolbox work)
- File 6: bound `.xml` model files (CobraPy/Python work)

- File 7: bound `.mat` model files (COBRA Toolbox work)
- File 8: `.html` MEMOTE report files
- File 9: `.json` Escher map files
- File 10: manifest file (`.xml`) describing the contents of the `.omex` file
- File 11: metadata file (`.rdf`) describing the authors

**Data S3**: Excel® workbook with fermentation data from first experiment, period 2

- Sheets 1-12: Time series data of $OD_{600}$, $g_{CDW}$, $H_2$, $CO_2$, $CH_4$, mmol/h and mmol/gCDW/h for all bioreactors
- Sheets 13-15: $OD_{600}$, $g_{CDW}$, dilution rates
- Sheets 16-23: Averaged steady-state time points for $H_2$, $CO_2$, $CH_4$, and biomass
- Sheet 24: Media salinity calculation
- Sheet 25: Dissolved $CO_2$ calculation
- Sheet 26: Carbon balance before gross measurement analysis/adjustment
- Sheet 27: Steady-state data after gross measurement analysis (which is used for modeling)

**Data S4**: Excel® workbook with FBA simulations with GEMs

- Sheets 1-3: FBA results of constraining the GEMs on test conditions for DH, ZZ, MM
- Sheets 4-6: FBA results of constraining GEMs on fermentation data (gross measurement adjusted) normal/Gimme Transcriptomics/Gimme Proteomics for DH, ZZ, MM

**Data S5**: Excel® workbook with fermentation data from second experiment

- Sheets 1-12: Time series data of $OD_{600}$, $g_{CDW}$, $H_2$, $CO_2$, $CH_4$, Na-formate mmol/h and mmol/gCDW/h for all bioreactors
- Sheets 13-15: $OD_{600}$, $g_{CDW}$, dilution rates
- Sheets 16-19: Averaged steady-state time points for $H_2$, $CO_2$, $CH_4$, and Na-formate

**Data S6**: GitHub repository with relevant scripts
https://github.com/isacasini/Casini_2022_GEM

- Script 1: Gross measurement error analysis `.ipynb`
- Script 2: Modeling 1 (CobraPy) used for model validation (tests the different conditions) `.ipynb`
- Script 3: Modeling 2 (CobraPy) used for model verification (constraints with fermentation data -adjusted with gross measurement error) `.ipynb`
- Script 4: Transcriptomics differential expression notebook in R `.ipynb`
- Script 5: Pan-proteome maker (w/ iBAQ script) `.ipynb`
- Script 6: MATLAB file to run FBAs through GIMME (COBRA Toolbox) `.mat`
